# Lys716 in the transmembrane domain of yeast mitofusin Fzo1 modulates anchoring and fusion

**DOI:** 10.1101/2023.09.22.559002

**Authors:** Raphaëlle Versini, Marc Baaden, Laetitia Cavellini, Mickaël M. Cohen, Antoine Taly, Patrick F.J. Fuchs

## Abstract

Outer mitochondrial membrane (OMM) fusion is an important process for the cell and organism survival, as its dysfunction is linked to neurodegenerative diseases and cancer. The OMM fusion is mediated by members of the dynamin-related protein (DRP) family, named mitofusins. The exact mechanism by which the mitofusins contribute to these diseases, as well as the exact molecular fusion mechanism mediated by mitofusin, remains elusive.

We have performed extensive multiscale molecular dynamics simulations using both coarse-grained and all-atom approaches to predict the dimerization of two transmembrane domain (TM) helices of the yeast mitofusin Fzo1. We identify specific residues, such as Lys716, that can modulate dimer stability. Comparison with a previous computational model reveals remarkable differences in helix crossing angles and interfacial contacts. Overall, however, the TM1-TM2 interface appears to be stable in the Martini and CHARMM force fields. Replica-exchange simulations further tune a detailed atomistic model, as confirmed by a remarkable agreement with an independent prediction of the Fzo1-Ugo1 complex by AlphaFold2. Functional implications, including a possible role of Lys716 that could affect membrane interactions during fusion, are suggested and consistent with experiments monitoring mitochondrial respiration of selected Fzo1 mutants.

## Introduction

Mitochondria form a complex network inside the cells, undergoing continuous fusion and fission events. These processes shape mitochondrial dynamics and are essential for the maintenance, function, distribution and inheritance of mitochondria. The morphology of the latter therefore respond to the ever-changing physiological changes of the cell [1].

Mitofusins are large GTPase transmembrane proteins involved in the tethering to and fusion of mitochondrial outer membranes (OM) [2]. The mitofusins Mfn1 and Mfn2 are found in mammals [3, 4]. Fzo1 (*Fuzzy Onion 1*) is the only mitofusin homolog in *Saccharomyces cerevisiae* [5]. The structure of Mfn1 was partially solved, but without its transmembrane domain [6, 7], and no solved structures are available for either Mfn2 or Fzo1. Mitochondrial inner membrane fusion and cristea organisation are mediated by human OPA1 (*Optic Atrophy 1*) [8] and yeast Mgm1 (*Mitochondrial Genome Maintenance 1*) [9].

Mitochondrial fusion dysfunction can cause neurodegenerative diseases such as Parkinson’s, Alzheimer’s, and Huntington’s [10, 11] as well as cancer [12, 13]. A number of studies have linked Mfn1 and Mfn2 to various cancers, including breast cancer [14], liver cancer [15], lung cancer [16], cervical cancer [17], and colon cancer [18]. They showed that their dysregulated expression was associated with increased cell proliferation, invasion, and resistance to chemotherapy. Regulation of mitofusin activity has been shown in preclinical studies to reduce cancer cell growth and spread. In addition, research has shown that mutations in Mfn2 trigger the development and progression of muscular dystrophies such as Charcot-Marie-Tooth type 2A, the most common form of axonal CMT disease[19, 20]. The precise mechanism by which mitofusins contribute to cancer or neurodegenerative disease, as well as the exact molecular fusion mechanism mediated by mitofusins, require further structural investigation.

From other membrane fusion machines such as the SNARE proteins, we know that anchoring in the membrane, transmembrane domain amino-acid composition, and dynamic properties are crucial elements that determine the interplay between the main players of the fusion process at different stages [21, 22, 23, 24]. With the help of a membrane-embedded structural model of Fzo1, such properties, the fusion itself, as well as its intermediate stages, could be studied by computational approaches, in particular molecular dynamics simulations. To this end, we are striving to build a complete structural model of Fzo1, including in particular the fusion-related transmembrane domains, for which we have very limited data.

Fzo1 has two heptad repeat domains (HRs) with coiled-coils on its N-terminal side: HRN (only in yeast) and HR1 flanking a GTPase domain. A third heptad repeat domain HR2 is located at the C-terminus [25, 26]. Some models of Fzo1 are based on the mitofusin-related bacterial dynamin-like protein (BDLP) [27, 28]. BDLP is involved in membrane remodelling and exists in two conformational states, a closed, compact version that transitions to an open, extended structure upon GTP binding. The latter structure can therefore bind lipid bilayers and results in self-accelerating polymerization leading to a coated lipid bilayer and strong curvature. However, BDLP has no transmembrane part, whereas Fzo1 is embedded in the mitochondrial OM via two transmembrane segments exposing N- and C-terminal parts to the cytosol and a loop to the intermembrane space [29]. The two transmembrane segments are putative *α*-helices called TM1 and TM2 [30].

We previously constructed an initial model of Fzo1 based on the closed compact conformation of BDLP, which was experimentally validated by mutation studies [30]. The opened extended structure of BDLP was used to create a second model [31, 32]. In the absence of a BDLP transmembrane template, the structure of TM1 and TM2 of Fzo1 was predicted using an *ab initio* method, namely the PREDDIMER [33] web server. Here we attempt a more comprehensive investigation of the possible membrane associations of the two helices to form a dimer using coarse-grained molecular dynamics simulations. Indeed, a new version of the popular Martini force field solved a previous bias favoring protein/protein association in membranes [34]. This new force field, Martini 3, therefore paved the way to more realistic studies of membrane protein associations[35]. In the case of Fzo1, the TM domain presents a particular challenge, as there is a basic conserved lysine residue in the middle of TM1. In addition to dimerization itself, we are investigating the extent to which the protonation state of this residue may affect dimer stability, membrane incorporation, and destabilization. Lysine protonation may be functionally relevant, as the literature has hypothesized specific roles for individual transmembrane domains in fusion and, in particular, charge-related effects. Specifically for the SNARE system, Lindau *et al.* postulated a mechanism in which the formation of fusion pores is triggered by the movement of the charged C-terminus of the transmembrane helix syb2 within the membrane [21].

Through extensive molecular simulations across coarse-grained and atomistic scales, we arrive at an improved model of the Fzo1 transmembrane domain that reveals new insights into the dimerization interface, key regulatory residues, and their potential influences on mitochondrial fusion.

## Materials and Methods

The transmembrane (TM) domain of Fzo1 contains two TM *α*-helices (called TM1 and TM2) connected by a loop, plus some flanking residues on the N-terminal part of TM1 and C-terminal part of TM2. In addition, TM1 possesses a lysine residue (Lys716) at its exact center, thus located in the middle of the membrane. This location therefore raises the question of its protonation state, knowing that the local pKa of lysine may go below 7.0 in very apolar environments [36, 37, 38].

The computational protocol used in this study consisted in two main parts. The first part used coarse-grained (CG) simulations for sampling the TM1/TM2 contacts and selecting the best possible TM1/TM2 dimer. Two CG force-fields (Martini 2 and Martini 3) were tested as well as the two protonation states of Lys716 (neutral or positively charged). In the second part, the best CG dimer was chosen, the loop between TM1 and TM2 added, the structure backmapped to an all-atom (AA) representation, followed by AA simulations to refine the whole Fzo1 TM domain. Everything is summarized in Figure 1. Below, we explain the details of each part. The GROMACS 2018.5 program was used to perform all molecular dynamics (MD) simulations [39].

**Figure 1.**
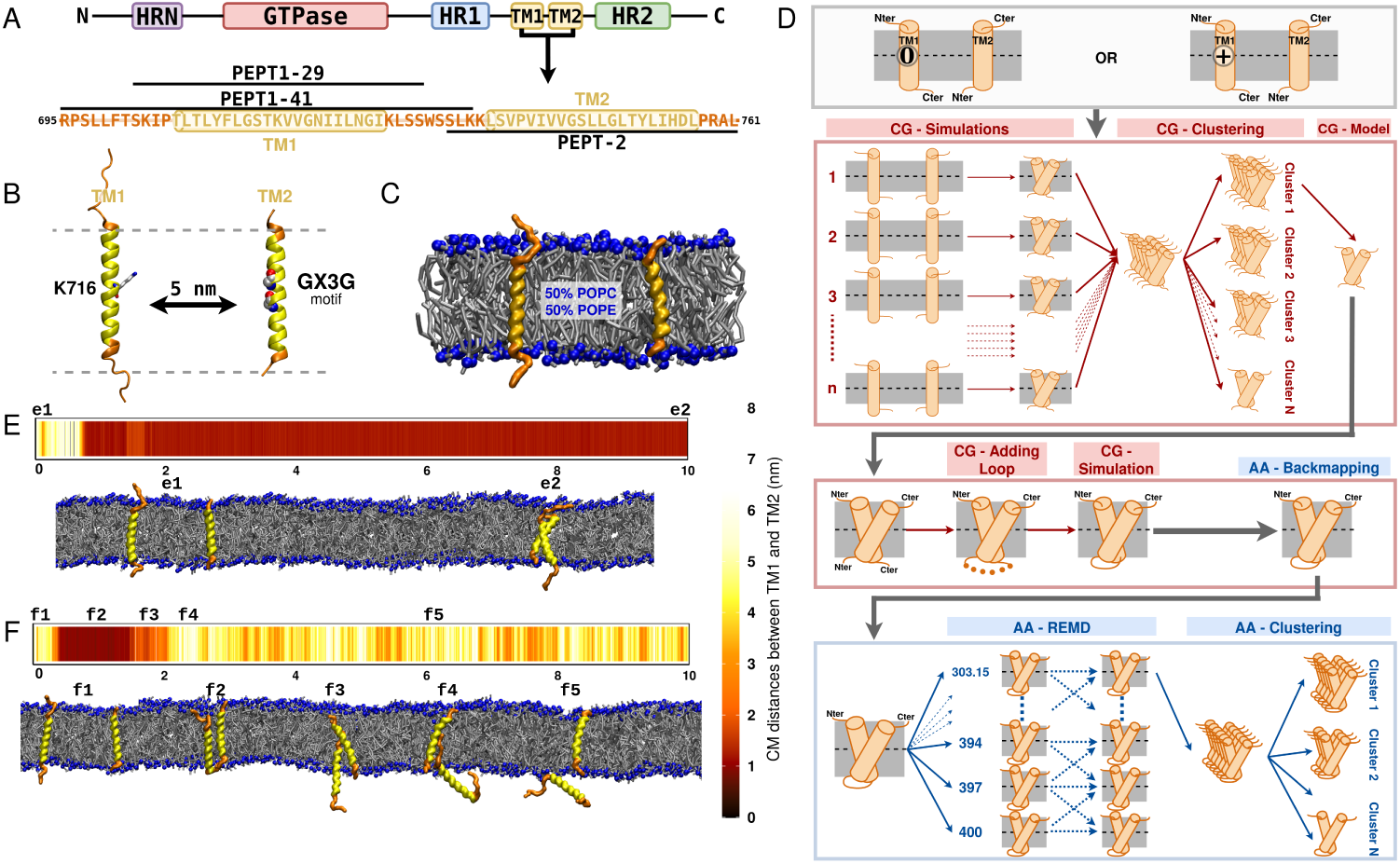
Protocol. (A) Sequence definition of the peptides PEPT1-29, PEPT1-41 and PEPT2. The TM domains are represented in yellow and the flanking residues are in orange. This color code is maintained for the snapshots in B,C,E,F. (B) Images of the all atom structures of the peptides PEPT1-41 and PEPT2. (C) Snapshot of the coarse-grained system. In blue are the phosphate groups, in silver are the lipids POPC and POPE. (D) A detailed scheme of the protocol. (E) Center of mass distances throughout the simulation, with snapshots of the beginning and end point. (F) Center of mass distances throughout the simulation, for a replica with TM1 expelled from the membrane. Snapshots throughout the simulations.

### Coarse-Grained conformational sampling

#### Peptide Definition

The *S. cerevisiae* Fzo1 sequence and domains were obtained from the UniProtKB database (Universal Protein Resource Knowledgebase [40], entry: P38297). The secondary structures of the protein were predicted using PSIPRED 4.0 [41, 42]. The positions of the transmembrane helices of Fzo1 were determined using TMHMM2.0 [43, 44]. Some peptides were defined (PEPT1 or PEPT2) which contained either TM1 or TM2 respectively, plus various flanking residues: one from Arg695 to Lys735 called PEPT1-41 (of 41 residues length), another from Ser702 to Ser730 called PEPT1-29 (of 29 residues length), and a last one from Ser733 to Leu761 called PEPT2 (of 29 residues length). PEPT1-29 and PEPT2 were used with Martini 2, whereas PEPT1-41 and PEPT2 were used with Martini 3. All sequences are shown in Figure 1A. The peptide 3D-structures were built using Basic Builder 1.0.2 on the Mobyle server (https://mobyle.rpbs.univ-paris-diderot.fr) [45]. The residues from Thr706 to Ile726 (TM1) and from Leu737 to Leu57 (TM2) were defined as a perfect *α*-helix and the flanking residues were defined as coil. This step gave a first all-atom 3D structure for each peptide, which was then used as an input to Martini and CHARMM-GUI tools described in the next section.

#### Box setup

When we started the work, Martini 2 was the main CG Martini force field and Martini 3 was in its beta version. However, the Martini 3 force field came out during the course of this work [34]. We thus decided to test both versions of the force-field, Martini 2 and Martini 3. We tested the two possible charge-states of Lys716 (positively charged or neutral), resulting in a total of four systems.

The Martini 2 systems were built using the CHARMM-GUI Membrane builder [46, 47, 48] with the Martini 2 coarse-grained (CG) force field [49, 50]. The two peptides were separately embedded in a mixed lipid bilayer of 50 palmitoyl-oleoyl-phosphatidylcholine (POPC) and 50 palmitoyl-oleoyl-phosphatidylethanolamine (POPE) (these two lipids are the most abundant in mitochodrion OM). Thus we had one box with PEPT1 (PEPT1-29 or PEPT1-41) and another one with PEPT2. About 40 water molecules per lipid were used as solvent for both PEPT1-29 and PEPT2, and about 50 water molecules per lipids were placed in the system of PEPT1-41. To the solvent were added 0.15M of NaCl for each system. The two systems were then assembled, in order to have the two peptides embedded in the same membrane at a distance of about 5.2 nm. In order to obtain a neutral Lys716 system, we replaced in the ITP file the Qd charged bead by a P1 neutral and polar bead.

Since the mapping between Martini 3 and Martini 2 is different, we could not convert the topologies directly. Instead, the Martini 3 systems were built from atomistic models of the peptides (using the same coordinates than for Martini 2) using the martinize2.py workflow module (see https://github. com/marrink-lab/vermouth-martinize). These Martini 3 structures of the peptides were then placed in lieu of the peptide structures in the Martini 2 system. In order to obtain a neutral Lys716 system, we replaced in the ITP file the SQ4p charged bead by a SN6d neutral and polar bead. As the mapping of the lipids did not change between the two versions of the force-field, we were able to use the membranes produced with Martini 2.

#### Simulation parameters

For all systems, we followed the protocol of CHARMM-GUI which consisted in two energy minimizations of 5000 steps, followed by an equilibration of 5 simulations within the NPT ensemble. The velocity-rescaling thermostat [51] at 303.15 K and the Berendsen barostat at 1 bar [52] were applied for a sequence of 5 simulations of 1 ns, 1 ns, 1 ns, 0.75 ns and 1 ns (with a timestep of 0.002 ps, 0.005 ps, 0.01 ps, 0.015 ps, 0.020 ps respectively). From the equilibrated system, a last step of 1 ns equilibration (time step 0.020 ps) with the same thermostat and barostat parameters was then used to redefine the velocities of the system (with a fixed seed). A production run of 10 *µ*s followed, at 303.15 K using the velocity-rescaling thermostat [51] (lipid, water and proteins coupled separately) and 1 bar using the Parrinello-Rahman barostat [53] (compressibility of 3.0 *×* 10*^−^*^4^*bar^−^*^1^). Pressure coupling was applied semi-isotropically. These final two steps were repeated multiple times, the seed being changed for each preceding equilibration step. A time step of 0.020 ps was used with the leapfrog integrator. Lennard-Jones interactions were cutoff at 1.1 nm. Bond lengths were constrained using the LINCS algorithm [54]. The reaction-field method [55] was used for evaluating electrostatic interactions, with a Coulomb distance cutoff of 1.1 nm, a relative dielectric constant of 15. The neighbor list was updated every 20 steps.

#### Clustering and model extraction

We ran as many simulations as necessary in order two obtain 25 trajectories ending in successful dimerization for each of the four conditions (two Martini versions and two protonation states). The unsuccessful trajectories were discarded. The successful ones were submitted to a conformational based clustering. The first 2 *µ*s of the productions were systematically ignored. All simulations were then concatenated (using 161 frames per simulation representing 8 *µ*s) resulting in a total of 4025 frames. This concatenated trajectory was used as input to the *gmx cluster* program of GROMACS. The GROMOS method [56] was used. Briefly, one first computes the pairwise RMSD matrix of TM1 and TM2 (RMSD of all pairs of conformations, considering backbone beads only) and then group conformations in different clusters. Within each cluster, any pair of conformations presents an RMSD below a cutoff that has to be chosen. This non-supervised method (the number of clusters is not fixed and is a result of the clustering) maximizes the size of the clusters. The cutoff was set to 0.3 nm in order to get 5 to 6 clusters totalizing 80 to 90 % of all conformations.

#### Trajectory analysis

The crossing angle between the TM domains was calculated as described by Chothia *et al.* [57] using an in-house script. Only the backbone beads of TM1 and TM2 were considered.

Contacts between TM1 and TM2 were studied using the python library MDAnalysis [58, 59]. The minimum distance between each residue pair was then plotted using the R library ggplot2 [60].

A principal component analysis (PCA) of the cartesian coordinates was carried out using the programs *gmx covar* and *gmx anaeig* of the GROMACS package. It was done on a combined trajectory of the dimerized structures of the 4 systems using the backbone beads of TM1 and TM2. The results are presented in terms of free energy projection against the two first principal components (PC1 and PC2) using a grid:

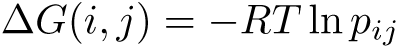

where *i* and *j* are indices of the grid points, *p_ij_*the probability of finding a conformation in grid point (*i, j*) and Δ*G*(*i, j*) the relative free energy of that grid point. The lowest free energy was set to 0 *kJmol^−^*^1^. 150 grid points were used for PC1 and PC2.

Lipid packing defects are small apolar areas of the membrane which are accessible to water. They were quantified using PackMem [61, 62]. A protrusion event is defined as one of the carbon atoms of a lipid tail bulging into the polar layer (or a backbone bead of a lipid tail), extending 0.1 nm above (or below, depending on the leaflet) its phosphorus atom [63, 64]. Protrusions were identified using an in-house script. Both analyses were performed separatly for each leaflet.

### All-Atom refinement of the model

The CHARMM36m force field for proteins [65] and CHARMM36 for lipids [66] were used for the remaining all-atom simulation described in this section.

#### System building and equilibration

Prior to all-atom simulations, the loop between TM1 and TM2 was reconstructed and sampled using CG simulations. We used position-restraints on TM1 and TM2 in order to maintain the contacts predicted in the previous phase. The details of this step can be found in the Supplementary Material. Briefly, the best loop conformation was chosen based on conformational clustering.

The whole TM domain including the loop was then backmapped using the CHARMM-GUI backmapping tool [67]. We reduced the box size in order to have 100 lipids located around the TM domains, as well as 40 water molecules per lipids and 0.15M of NaCl. In total, the system consisted in 26144 atoms. The preceding step introduced a layer of vacuum on the box edges. Using 8 minimizations of 5000 steps, we progressively shrinked the box size to get rid of this vacuum layer and recover periodic boundary conditions. The system was then submitted to a sequence of 6 equilibrations of 125 ps, 125 ps, 2 ns, 2 ns, 2 ns, and 2 ns (with a timestep of 0.001 ps, 0.001 ps, 0.001 ps, 0.002 ps, 0.002 ps, 0.002 ps respectively) in which we progressively released the position-restraints on the protein. The first two equilibrations were performed within the NVT ensemble, with the Berendsen thermostat, and the following simulations were performed within the NPT ensemble, with the Berendsen thermostat and barostat. The temperature was maintained at 303.15 K and the pressure at 1 bar. Pressure was applied semi-isotropically. Electrostatic interactions were calculated with the particle-mesh-Ewald (PME) method [68, 69], with a real-space cutoff of 1 nm. Van der Waals interactions were computed using a Lennard-Jones force-switching function over 10 to 12 Å. Bond lengths were constrained using the LINCS algorithm [70]. Water molecules were kept rigid with the SETTLE algorithm [71].

#### Temperature replica-exchange molecular dynamics

In order to explore efficiently the conformational landscape of the TM domain, we used temperature replica-exchange molecular dynamics (T-REMD)[72]. The replica temperatures were predicted using the webserver https://virtualchemistry.org/remd-temperature-generator/ [73] by setting the exchange probability to 0.2 and the temperature range between 303.15 and 399.83 K. The upper limit of nearly 400 K was chosen so that conformational sampling of the protein was more efficient, but the membrane stayed intact (if heated too much, it can explode). In total, we obtained 38 replica.

Each replica was equilibrated at the chosen temperature for 1 ns (time step 0.002 ps) using the Berendsen thermostat and barostat (at 1 bar) with different starting velocities. The production run of 500 ns followed with the velocity-rescaling thermostat [51] (lipid, water and proteins coupled separately) and the Parrinello-Rahman barostat at 1 bar [53] (compressibility of 4.5 *×* 10*^−^*^5^*bar^−^*^1^). Exchanges between neighboring replicas were attempted every 10 ps. The other settings were identical to those described in the previous section.

#### Trajectory analysis

All the analyses were performed either based on individual replica that diffuse in temperature, or on the conformations of the bottom temperature (303.15 K). On the latter, crossing-angles (using helical C*α* atoms only), conformational clustering (using a cutoff of 0.2 nm and a distance between the TM1 and TM2 center of mass below 2 nm) and contact-maps were calculated using the same programs as for the CG simulations. The cutoff of 0.2 nm used for the clustering was chosen so that each cluster roughly coincided with the group of points obtained in the PCA analysis below.

A PCA was carried out on a trajectory combining the REMD structures used for the clustering and the Martini 3 neutral Lys716 structures. To do this, the all-atom conformations were converted to a CG representation, and concatenated to the Martini 3 structures for the covariance matrix calculation. Only the backbone beads of TM1 and TM2 were considered.

A conformational clustering was performed on the C*α* atoms of TM1 and TM2 using the same algorithm and tool as for CG simulations (cutoff of 0.2 nm). The central structure of the first and third clusters were then extracted. We selected these two structures because of their positioning within the deepest energy wells identified by the previous PCA.

Hydrogen bonds were calculated using *gmx hbond* from GROMACS 2018.5. Contacts within a distance cut-off of 0.35 nm and up to 30 degree off-axis angle were considered.

### AlphaFold2 predictions

To predict the structures of Fzo1, we used Alphafold version 2.2 [74, 75], and Colabfold 1.3.0 [76], both monomer and multimer versions. The provided sequences of Fzo1 (from residues 491 to 813, total length of 323 residues, Uniprot: P38297) and Ugo1 (Uniprot: Q03327) were taken from Uniprot [40].

### Plasmids,yeast strains and growth conditions

Mutations on K716 (plasmids MC430, MC437, MC587 and MC589, Table1) were generated by PCR using QuikChange Lightning Site-Directed Mutagenesis Kit (Agilent Technologies 210518, Santa Clara, California, USA). The QuikChange Primer Design Program is available online at https://www.agilent.com/store/primerDesignProgram.jsp and was used to design mutagenic primers based on the Fzo1 sequence. Standard methods were used for growth, transformation and genetic manipulation of S. cerevisiae. Minimal synthetic media [Difco yeast nitrogen base 291940 (Voigt Global Distribution, Inc., Lawrence, Kansas, USA), and drop-out solution] supplemented with 2% dextrose (SD) or 2% glycerol (SG) were prepared as previously described in Sherman et al, J. Methods in Yeast Genetics (1986)[77]. All experiments were performed with the fzo1D shuffle strain MCY571 where mitochondrial fusion efficiency is maintained by the pRS416-FZO1 shuffle plasmid carrying a copy of the Fzo1 wild-type gene and the URA3 selection marker. This strain was transformed with the pRS314 FZO1 plasmids described in Table 1 carrying the TRP1 selection marker and resulting transformants were plated on SD selective media lacking uracil and tryptophan. Colonies were isolated on SD selective media and replica-plated on 5’-fluoroorotic acid (5’-FOA) plates to cure the strains from the FZO1 shuffling plasmid.

**Table 1.**
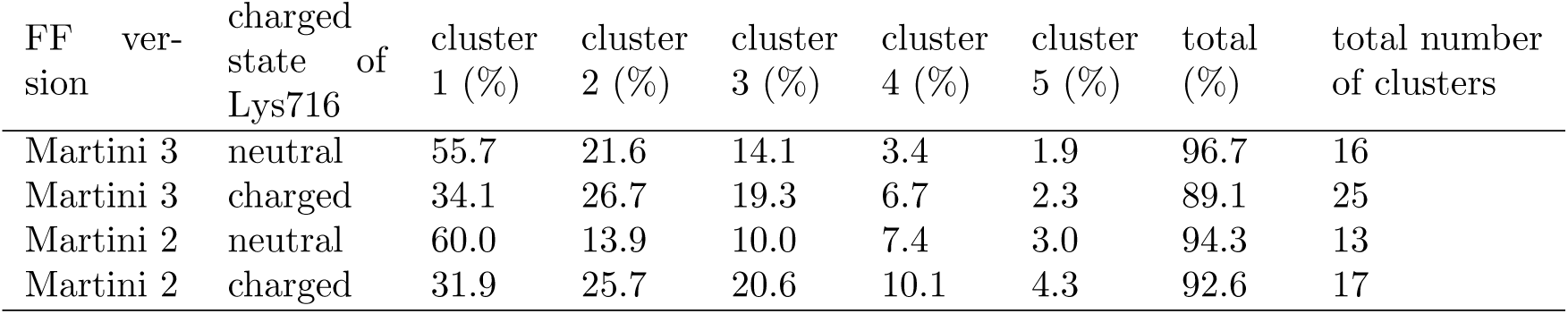
Plasmids used in this study.

### Spot assays

Cultures grown overnight in SD selective media lacking tryptophan were pelleted, resuspended at OD600=1 and serially diluted (1:10) four times in water. Five microliters of the dilutions were spotted onto SD and SG selective plates and grown for 2 days at 23 °C, 30 °C or 37 °C.

### Protein extracts and immunoblotting

Cells grown in SD selective media lacking tryptophan were collected during the exponential growth phase (OD600 = 0.7–1). Total protein extracts were prepared using the NaOH/trichloroacetic acid (TCA) lysis technique[78]. Proteins were separated on 8% SDS-PAGE gels and transferred to nitrocellulose membranes (Amersham Protran 0,45 µm 10600002; GE Healthcare, Little Chalfont, United Kingdom). The primary antibodies used for immunoblotting were monoclonal anti-Pgk1 (Abcam ab113687, Cambridge,United Kingdom) and polyclonal anti-Fzo1 (generated by Covalab, Bron, France). Primary antibodies were detected using horseradish peroxidase-conjugated secondary anti-mouse or anti-rabbit antibodies (HRP, Sigma-Aldrich 12-348 and A5278, Saint-Louis, Missouri, USA), followed by incubation with a Clarity Western ECL Substrate (Bio-Rad 1705060, Hercules, California, USA). Images of the immunoblots were acquired using a Gel DocTM XR+ (Bio-Rad) and analysed using the Image Lab 3.0.1 software (Bio-Rad).

## Data availability

R-markdown notebook and Zenodo archive.

## Results

### The association of transmembrane helices is reproducible with two coarsegrained force fields

Our aim was to predict the structure of the transmembrane domain of Fzo1 in a 1:1 POPC-POPE bilayer. This lipid composition was chosen considering the two most abundant phospholipids in the outer mitochondrial membrane and was the same as in our previous study [30]. To evaluate the possible associations between the two helices TM1 and TM2 in coarse-grained (CG) simulations, the domain was cut into two parts and some peptides were defined containing either TM1 or TM2: PEPT1-29, PEPT1-41, and PEPT2. Each sequence contains also some flanking residues on both the N- and C-terminal sides (see Figure 1A). In the simulations performed with the Martini 2 force field, PEPT1-29 and PEPT2 were used. However, in the simulations performed with the Martini 3 force field, this short version of the TM1-containing peptide, PEPT1-29, was observed to exit the membrane, preventing dimerization events. Inspired by a recent study of Martini 3 dimerization of TM-helices [35], we replaced the short sequence with the longer PEPT1-41 variant, which contains additional flanking residues at both the N- and C-termini, effectively reducing the number of ejections. The longer TM1 sequence allowed us to generate 25 successful dimerizations (for each protonation state of Lys716) for subsequent analyses. Because the simulations with Martini 2 and Martini 3 produced quite similar results in terms of dimer associations, and because Martini 3 addresses a number of shortcomings of Martini 2, such as exaggerated protein-protein aggregation, we decided to present only the Martini 3-based results in the remaining sections of this article. The corresponding Martini 2 results can be found in supplementary Figure 9.

### Protonation of Lys716 interferes with the formation of a stable TM1-TM2 dimer

The simulations were started with a distance between the two peptides of about 5.2 nm. Most of the time, spontaneous and irreversible dimerization was observed (Figure 1 e1-e2 and f1-f2), but sometimes the peptide containing TM1 was ejected from the membrane (Figure 1 f3-f4-f5). All ejections occurred on the intermembrane side, which corresponds to the lower leaflet in our simulations. When dimerization occurred, we observed that the distance between the two peptides stabilized at 1.5 nm. This event generally occurred within the first 2 *µ*s in each replicate (supplementary Figure 6A). In the Martini 2 simulations, dimerization was irreversible in all runs (supplementary Figure 6A). In contrast, in a considerable number of Martini 3 simulations, TM1 ejection occurred, which affected the dimerization process. We therefore had to run up to 63 simulations with Martini 3 to obtain the targeted set of 25 trajectories that ended with a stable dimer (supplementary Figure 6A and 6B). For charged Lys716, we observed almost 5 times more ejections (38 ejections versus 25 dimerizations) than for the neutral version (8 ejections versus 25 dimerizations) (see supplementary Figure 6). The lower propensity for dimerization observed with Martini 3 when Lys716 is charged underscores the greater compatibility of a neutral form for self-association within the hydrophobic environment of the membrane.

### Protonation of Lys716 acts as a switch for remodeling the TM1-TM2 interface

Next, we analyzed the TM1-TM2 contacts that resulted from the CG simulations after dimerization. To identify the most important dimer conformations from the two sets of 25 trajectories, we clustered the dimers using the GROMOS method [56]. This clustering was based on the pairwise RMSD matrix of the backbone beads of each pair of dimerized TM domain conformations. In the Martini 3 simulations with a neutral Lys716, clustering resulted in 5 clusters representing 96% of the conformations, with the first 3 clusters corresponding to 91.4% of the structures. However, when Lys716 was charged, we found that the first cluster was smaller, while the others were larger. Also, the total number of clusters was larger with charged Lys716. This observation is an indication of a greater conformational diversity. The same trend was observed in the Martini 2 simulations (Table 3, Supplementary Figure 9).

**Table 2.** Saccharomyces cerevisiae strains used in this study.

| Name | Genotype | Reference |
| --- | --- | --- |
| FZO1 (MCY571) | MATα ura3-1 trp1-1 leu2-3,112 his3-11,15 can1-100<br>fzo1Δ::LEU2 pRS416-FZO1 | [30] |
| fzo1Δ (MCY620) | MATα ura3-1 trp1-1 leu2-3,112 his3-11,15 can1-100<br>fzo1Δ::LEU2 pRS314 | This study |
| FZO1 (MCY1261) | MATα ura3-1 trp1-1 leu2-3,112 his3-11,15 can1-100<br>fzo1Δ::LEU2 pRS314-FZO1 | This study |
| FZO1-K716R (MCY1402) | MATα ura3-1 trp1-1 leu2-3,112 his3-11,15 can1-100<br>fzo1Δ::LEU2 pRS314-FZO1-K716R | This study |
| FZO1-K716S (MCY1425) | MATα ura3-1 trp1-1 leu2-3,112 his3-11,15 can1-100<br>fzo1Δ::LEU2 pRS314-FZO1-K716S | This study |
| FZO1-K716V (MCY2258) | MATα ura3-1 trp1-1 leu2-3,112 his3-11,15 can1-100<br>fzo1Δ::LEU2 pRS314-FZO1-K716V | This study |
| FZO1-K716I (MCY2259) | MATα ura3-1 trp1-1 leu2-3,112 his3-11,15 can1-100<br>fzo1Δ::LEU2 pRS314-FZO1-K716I | This study |

**Table 3.**
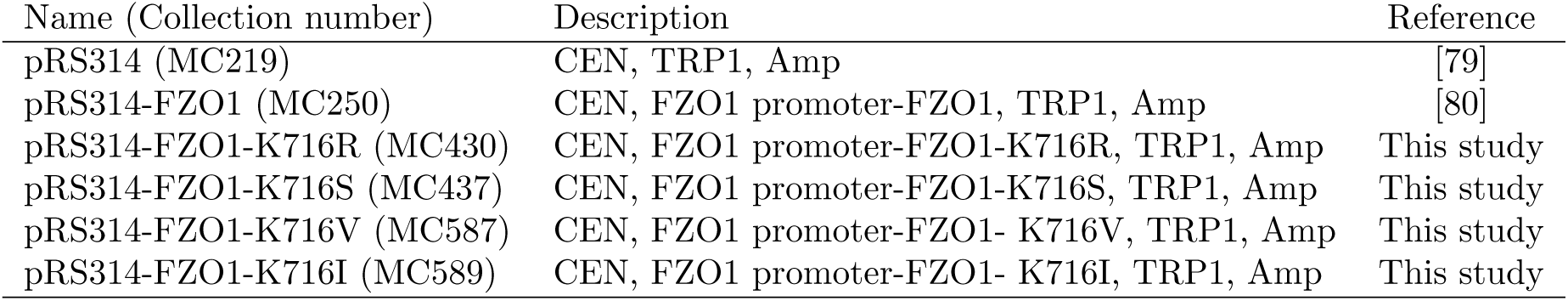
Cluster populations from the CG Martini simulations. Shown is the fraction of the 5 first clusters, the sum of these 5 fractions and the total number of clusters.

The interactions between the two TM helices can be observed conveniently by plotting the position of the center of mass of TM1 relative to TM2. All structures used for the clustering analysis were fitted to the backbone beads of TM2, and the positions of the center of mass of TM1 were then plotted in Figure 2A,B. This plot shows the position of TM1 around TM2, which is fixed in the center. Overall, this analysis reveals that the charge state of Lys716 controls the TM1-TM2 association. For charged Lys716, TM1 contacts TM2 on the left and top (Figure 2B), whereas for neutral Lys716, TM1 contacts TM2 on the right and top (Figure 2A). TM-TM contacts are therefore completely different depending on the charge state of Lys716.

**Figure 2.**
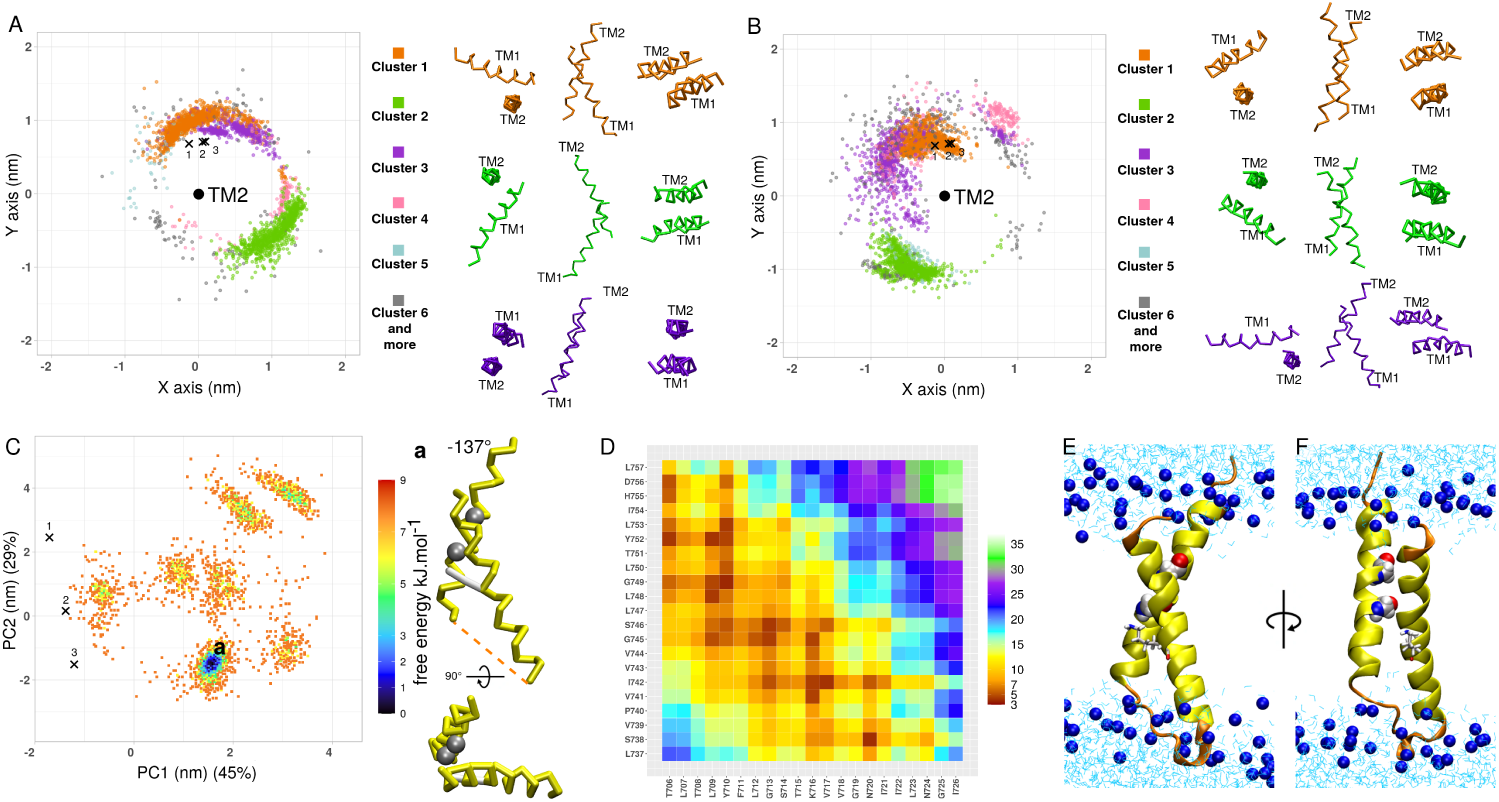
TM1-TM2 contacts determined using Martini 3 and best model extraction. (A,B) Positions of the center of mass of TM1 around the center of mass of TM2 (midpoint) for neutral (A) and charged (B) Lys716. The coloring of TM1 positions is based on the cluster to which the structure belongs to. Shown on the right side of each plot are the central structures of the first three clusters, colored similarly. (C) Free energy projection on the first two principal components of a PCA analysis (neutral Lys716 system). This PCA was performed on a concatenated trajectory containing all CG simulations (Martini 2 and 3, neutral and charged Lys716). The crosses 1, 2 and 3 correspond to the 3 dimers obtained with PREDDIMER in ref. [30]. Shown on the right are two snapshots of the best model (labeled “a” and corresponding to the center of cluster 1 in panel A and to the deepest free energy well), together with its crossing angle. The glycines of the GX_3_G motif are shown in van der Waals representation. Lys716 is shown in stick representation. (D) Contact map of the best model. (E,F) Best model back-mapped to an all-atom representation with the loop between TM1 and TM2 added. The protein is shown in a cartoon representation. TM1 and TM2 are shown in yellow and the flanking residues (with the loop) are shown in orange. Lys716 is shown in stick representation, and the GX_3_G motifs are shown in van der Waals representation. The phosphorus atoms of the lipids POPC and POPE are shown as blue spheres, and the water molecules are shown as cyan sticks. The loop was added according to the procedure described in the supplementary material.

### The TM domain energy landscape quantifies Lys716-mediated dimer (de)stabilization

A principal component analysis (PCA) was performed on the dimerized structures from the 4 sets of simulations. Figure 2C shows the projection of each conformation onto the first two principal components, in terms of free energy, for neutral Lys716 with Martini 3. For charged Lys716 (as well as for Martini 2 simulations), the PCA plots are shown in supplementary Figure 9. For neutral Lys716, there is one main free energy well (Figure 2C), whereas there are several shallow wells when it is charged (supplementary Figure 9). This observation is consistent with the clustering results (Table 3). The charged residue promotes the exploration of a greater variety of conformations. We compared these results with dimers previously obtained in Ref. [30] predicted with PREDDIMER [33]. It can be observed that none of the PREDDIMER predictions match the first 5 clusters or the deepest wells (Figure 2A, 2B, 2C).

For the neutral Lys716 system, the deepest free energy well (Figure 2C) coincides exactly with the center of the first (largest) cluster (Figure 2A). Therefore, we decided to choose this latter structure as our best CG model. We extracted it for further analyses and conformational sampling at the allatom level (see below). This best model shows a compact structure characterized by a crossing angle of *−*137.4*^◦^* (Figure 2C) and it is a left-handed antiparallel dimer. The GX_3_G motif contained in TM2 is involved in the interaction with TM1 (Figure 2C), consistent with the literature [81]. However, this motif is usually observed in right-handed dimers [82, 83, 84], which is not the preferred arrangement here.

To have a better idea of the contacts, Figure 2D shows a contact-map of this best CG structure, and Figures 2E and 2F show a back-mapping of it to an all-atom representation. The part of each helix involved in the interaction ranges from T706 to N720 for TM1 and S738 to L753 for TM2. The N-terminal part of TM1 interacts mainly with the C-terminal part of TM2 (Figure 2B).

### Charged Lys716 destabilizes the membrane

The presence of a Lys residue in the middle of a TM helix raises the question of its putative role in the fusion process. Here, we performed CG simulations of the dimerization of TM1 and TM2 (as well as some flanking residues) with charged or neutral Lys716, allowing us to assess the effects of the dimers on the surrounding membrane environment. Thus, we analyzed two parameters associated with the onset of hemifusion, the fusion of the outer layers of each membrane: (i) lipid packing defects, which quantify the hydrophobic membrane surface in contact with the solvent [61, 62] and (ii) lipid tail protrusions, defined as the appearance of a carbon atom (or a coarse-grained bead belonging to a lipid tail) protruding above the level of the phosphate group [63, 64]. When such a protrusion occurs, the hydrophobic lipid tail is assumed to be in contact with the solvent. Thus, both parameters indicate the probability that the hydrophobic tails are exposed to the solvent, which is a prerequisite for hemifusion [85, 86, 87].

For lipid packing defects, the *π* constant provides information on the occurrence and size of such defects, with a higher value indicating more frequent and larger defects. Interestingly, the *π* constants are higher when the TM domains are present compared to pure membranes (supplementary Figure 7A). The constant for charged Lys716 is slightly higher, which suggests more packing defects, than that for neutral Lys716. The effect is however too modest to draw a definitive conclusion for this parameter. As for the protrusions, the presence of the protein favors their occurrence (Supplementary Figures 7B), which is consistent with the analysis of packing defects. In addition, an increased occurrence of protrusion events is observed when Lys716 is charged, indicating a destabilizing effect of the charge on the membrane. In summary, the route to hemifusion is favored not only by the presence of the TM domain, but the charge state of Lys716 also matters.

### Replica exchange MD provides a refined atomistic model and verifies stable TM assembly

To fine-tune a model of the whole TM domain, we performed all-atom simulations (AA) using an enhanced sampling technique, temperature replica-exchange molecular dynamics (T-REMD). As a starting structure, we chose the best dimer of the Martini 3 simulations obtained with neutral Lys716, as described above and in Figure 2. The missing loop was added and sampled as described in the supplementary methods. Finally, the system was back-mapped to an AA representation (see Figure 2E and 2F) and used as the initial structure for a T-REMD simulation of 500 ns. In total, 38 replica were simulated ranging from 303.15 K to 400 K. This REMD protocol allowed us to test the robustness of the model by exposing it to high temperatures while maintaining a physical Boltzmann distribution at each temperature.

We first analyzed the behavior of each replica as it propagated through temperature space. Of the 38 replicas, 19 showed persistent interactions between the two TM helices (center of mass / center of mass distance below 2 nm) throughout the simulation, 12 showed TM dissociations, while 7 showed TM dissociations and re-associations (Supplementary Figure 10A). Importantly, the C*α* RMSD of TM1 and TM2 showed minimal structural changes for each replica when the two TM regions maintained their interaction during the simulation (Supplementary Figure 10B), indicating stability and robustness of the original structure.

Next, we focused on the ensemble of structures at the lowest temperature (303.15 K) (Figure 3). In total, 79.2 % of the conformations presented stable TM1 / TM2 contacts. In addition, all conformations showed stability of their secondary structures in the two TM helices, even those which displayed dissociation (Supplementary Figure 11). Figure 3A shows the position of TM1 around TM2 fixed at the center, similar to Figure 2A for CG simulations, in terms of free energy. The conformations of the bottom temperature quickly left the area of the starting structure (represented by a cross), and populated mainly the right side of TM2, as well as its upper side but in a less pronounced way. The Martini 3 simulations had captured the right and upper parts overall, but not the exact upper right region found here.

**Figure 3.**
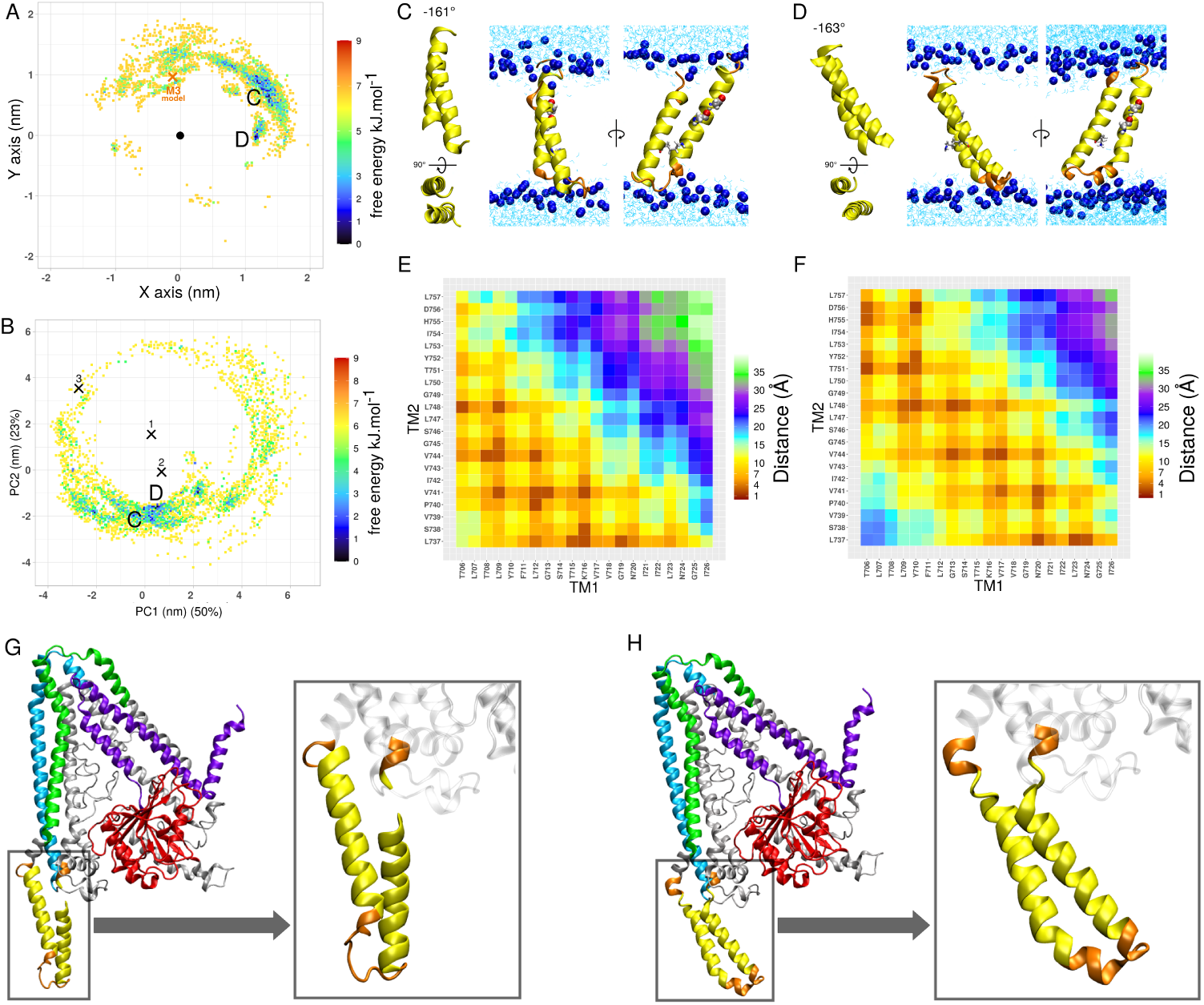
TM domain prediction based on REMD simulations. (A) Positions of the center of mass of TM1 around the center of mass of TM2 (center point) colored as function of the free energy. The orange cross is the best CG model obtained with Martini 3 (in Figure 2) used as a starting point of the REMD. (B) Free energy projection on the first two principal components of a PCA analysis. This PCA was performed on a concatenated trajectory containing Martini 3 (neutral K716) and all-atom (bottom temperature) conformations. On the plot, only the all-atom data are shown. The crosses 1, 2 and 3 correspond to the 3 dimers obtained with PREDDIMER in ref. [30]. The structures selected for both (A, B) have a TM1-TM2 COM-COM (center-of-mass) distance of less than 2 nm. (C,D) Center structure of the first and third cluster respectively, for a cutoff of 0.2 nm. These clusters roughly correspond to the 2 main free energy wells (bluest parts) seen in (A,B) and labeled C and D. The TM domains and the flanking residues are in cartoon representation. TM1 and TM2 are in yellow, and the flanking residues (with the loop) are in orange. The glycines of the GX_3_G motif are shown in van der Waals representation. The Lys716 is shown in stick representation. (E,F) Contact map of the models (C,D) respectively. (G,H) Addition of respectively (C) and (D) to the model of Fzo1 built by De Vecchis *et al.* in 2017 [30]. TM1 and TM2 are seen in yellow, the other residues used for the REMD are seen in orange, HRN is in purple, HR1 in blue, HR2 in green and the domain GTPase is in red. The residues that do not belong to a specific domain are in light grey. The second image is the TM portion, zoomed in. The residues before Ser702 and after Lys761 are represented in transparent gray.

In the following, we consider only the conformations with TM1 / TM2 in contact. From this ensemble of structures, we extracted the two best models of the entire TM domain from the two wells with the lowest free energy, as shown in Figure 3A-F. The two wells are labeled C and D in panels 3A and 3B, whose structures are shown in panels 3C and 3D. In both models C and D, TM1 touches TM2 in the lower part (the most likely contacts in Figure 3E-F are below the diagonal of the contact map), especially in model C. Both models C and D were then compared with the overall Fzo1 model that we proposed in 2017 [30] to check which model fits better (Figure 3G-H). As can be seen, model D (Figure 3D/H) fits better with the overall model of Fzo1 because the TM domain can be connected to the rest of the model at both the N- and C-terminal sides. We therefore selected model D as our best all-atom model. However, it is important to keep in mind that the TM contacts have a certain degree of flexibility, so that the alternative conformation C cannot be completely excluded.

We also analysed the promiscuous stabilizing interactions between TM1 and TM2 (Figure 3E-F). While we observed a similar set of residues involved in the interactions in TM1, there is a clear shift in the interactions involving TM2 (Figure 3). Instead of Gly745 and Gly749 of the GX_3_G motif, residues Val741 and Leu748 were the most involved in the interactions. We also found that among the structures located within the energy wells (Figure 3), Thr706, Leu709, Gly713, Lys716 from TM1 and Leu737, Val741, Val744, Leu748 from TM2 were the most promiscuous residues.

In terms of hydrogen bonds, we found some between the side chains of Lys716 and Ser738 with 8% of persistence. However, the latter residue frequently formed hydrogen bonds with the side chain of the TM1 residue Asn720 (12% of persistence) as well as with the residues of the loop from Lys726 to Lys735 (both side chains and backbone with 7% of persistence) and most frequently with the solvent (21% of persistence).

### Experiments show that the polar residue 716 is crucial for mitochondrial respiration in yeast

We sought to obtain experimental confirmation of our theoretic observations on residue K716. We reasoned that mutating this Lysine into apolar residues (I or V) should impact mitochondrial fusion efficiency while mutation into polar (S) or charged (R) residues should have no effect. To test this prediction, we took advantage of the established link between mitochondrial fusion efficiency and respiratory growth of Saccharomyces cerevisiae. Depending on the carbon source provided in their media, yeast cells grow through either fermentation or respiration. Dextrose is a fermentative carbon source that inhibits respiration whereas Glycerol is a fully respiratory carbon entry [88, 89]. Consequently, since mitochondrial fusion is essential for respiration, its inhibition abolishes yeast growth in Glycerol-containing media [90]. WT, fzo1Δ and cells expressing distinct versions of Fzo1 mutated in K716 as the sole source of Fzo1 (Fig. 4a) were thus subjected to serial dilutions growth assays on Dextrose or Glycerol-containing media at 23, 30 or 37°C (Fig. 4b). All mutants were expressed at levels comparable to WT Fzo1 (Fig. 4A). However, respiratory growth of FZO1 K716I and FZO1 K716V cells was abolished at all temperatures, similar to fzo1Δ cells. In contrast, respiratory growth of FZO1 K716S and FZO1 K716R cells was not affected, similar to WT cells. These results indicate that mutation of Lys716 to an apolar residue (K716I or K716V) impairs respiration likely because of an inhibition of mitochondrial fusion. Moreover, mutation of Lys716 to an Arg (K716R), the main alternative in Mammalian mitofusins, or Mutation of Lys716 to a polar residue (K716S) has no effect on respiration. In this context, the positive charge of Arg or Lys does not seem to be mandatory, but the residue polarity of Lys, Arg or Ser at position 716 would be essential for mitochondrial fusion.

**Figure 4.**
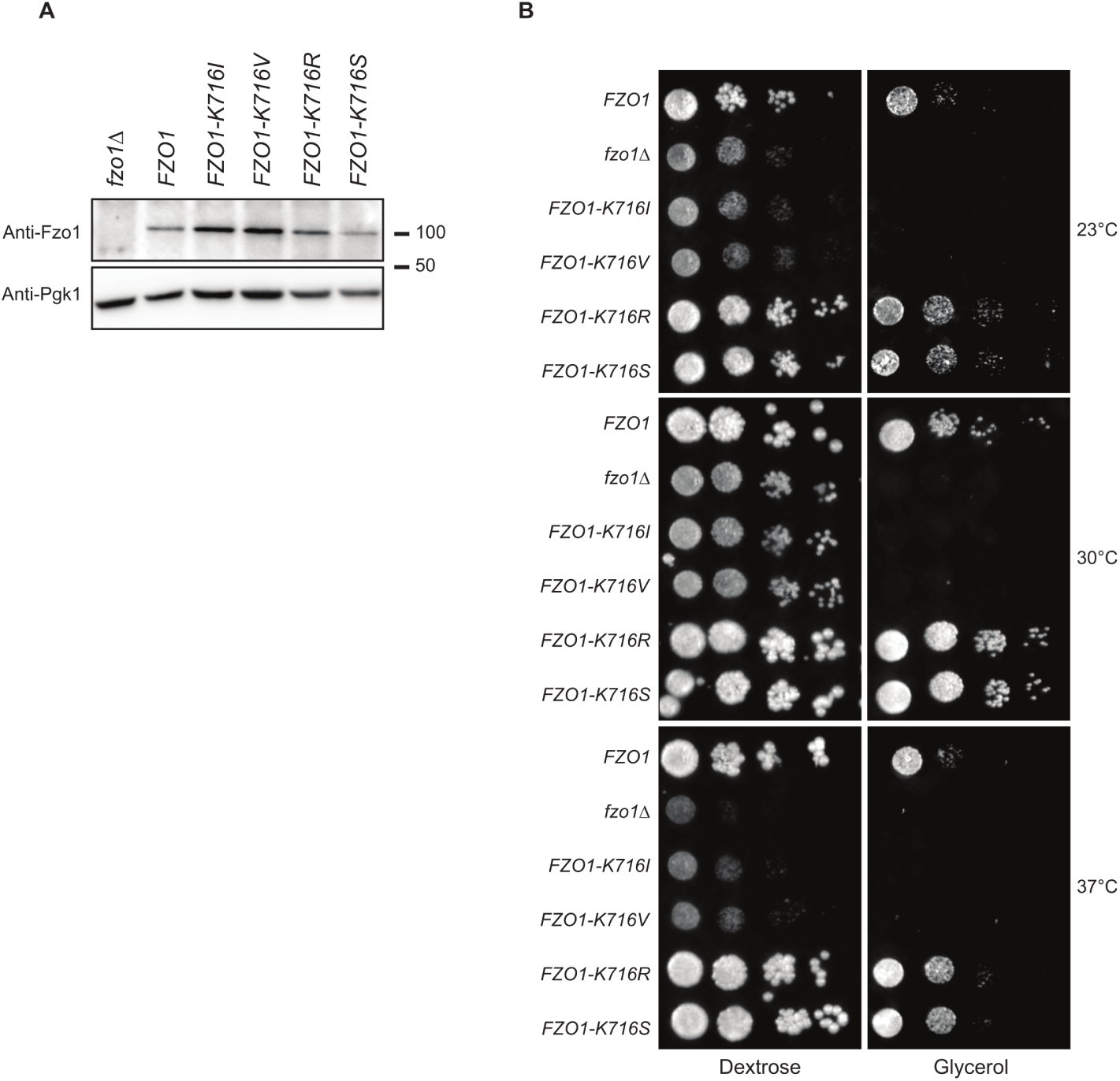
The polarity of K716 is essential for Fzo1 function. A) Anti-Fzo1 and anti-Pgk1 immunoblots of total protein extracts prepared from, fzo1Δ shuffle strains transformed with an empty plasmid or plasmids expressing FZO1 WT or mutated in K716. B) Dextrose and glycerol growth spot assay with strains used in A domain contacts. Next, we explain how our extensive multiscale simulations have led to a new structural model that significantly improves on previous computational predictions for the Fzo1 TM domain. We analyze the areas where our model deviates and discuss on the possible reasons for the increased accuracy. An independent prediction of the Fzo1-Ugo1 complex from AlphaFold2 is used to externally validate our new TM domain fold. We examine the remarkable agreement between these completely different approaches. We then discuss the functional implications of our refined structure, including how specific residues may be actively involved in the membrane fusion process. Our model sheds light on how the properties of the TM domain may facilitate the hemifusion and full fusion stages. Finally, we compare the results with other studies of dimerization, which give a consistent picture of such processes when examined with coarse-grained MD simulations.

## Discussion

Our extensive simulations provide new insights into the structure and dynamics of the transmembrane domain of Fzo1. Comparison with previous predictions reveals important differences, while independent validation suggests that the refined model accurately reproduces the folding of the TM domain. In this discussion, we first examine the robustness of the predicted TM1-TM2 interface in different simulation methods. Comparison of coarse-grained and all-atom approaches shows the robustness of the main TM

### The TM1-TM2 interface is robust across force fields and levels of representation

To investigate the possible association of TM1 and TM2, we generated twenty-five 10 *µ*s coarse-grained (CG) MD trajectories showing dimerization. Although the TM1-TM2 biological construct is connected by a loop in the intermembrane space, we intentionally cut the TM domain into two peptides so that dimerization was freely driven by TM1-TM2 contacts within the membrane. This assembly approach can be considered equivalent to docking, but confined within an explicit membrane environment. Because we used the CG Martini force fields [49, 50, 34] with smoother energy surfaces than a fully atomistic model, we were able to benefit from accelerated kinetics and thus efficiently sample the different modes of interaction of the two TM helices. As a result, we obtained robust statistics on TM1-TM2 contacts. Our protocol was first tested with the Martini 2 force field (supplementary Figure 9). During the progress of this work the Martini 3 force field was released, which encouraged us to use it as well. Interestingly, the Martini 2 simulations yielded a very similar dimerization pattern compared to Martini 3, albeit with a stronger association, which is a known artefact of Martini 2 [34]. Conformational clustering showed the same trends for both force fields, i.e., the charged Lys716 system has a larger number of clusters, and the first cluster of the simulations with neutral Lys716 accounts for more than 50% of the sampled structures. Experimentally, some ^2^H experiments were performed on a TM helix containing a Lys in its center [37]. A charged Lys was leading to a multistate behavior in terms of peptide orientation in the membrane (tilt and azimuthal rotation), whereas a neutral Lys gave a single orientation. In our CG simulations we have two TM helices dimerized, but we find the same kind of behavior since we get a single free energy well for neutral Lys716, and multiple shallow ones for the charged version.

When the model was subjected to all-atom simulations (AA) using an enhanced sampling method (T-REMD), we observed a slight shift in the most frequent interactions between the two TM helices. Lamprakis *et al.* showed that Martini 3 does not always favor the experimentally solved interface of interactions between TM domains [91] and recommended that a refinement procedure be used. Nevertheless, the interfaces of the interactions involving TM1 are overall the same in the Martini 3 and CHARMM36 refinements. The change to a detailed AA representation affected mainly the interactions involving TM2, while in the overall model, the interface of the interactions and the crossing angle were only slightly affected.

### How can we interpret TM1 expulsion from the membrane with Martini 3?

Next to experiments of biophysics or cellular biology, CG simulations with Martini 2 or 3 have been widely used to predict TM-TM helix dimerization (or higher order oligomerization), for example in recent works [92, 93, 94, 95]. Both force field versions are able to predict TM-TM interfaces, but Martini 2 has a tendency to overaggregate TM segments. Considerable efforts have thus been put in the development of Martini 3 to solve this issue [34]. This new version was recently tested on many known TM-TM dimers with success [35], but the authors highlighted the ability of Martini 3 to sample alternative conformations as well as the importance of flanking residues (around TM helices). Subsequent studies found the possible ejection of TM helices with Martini 3 [95, 96], ending adsorbed on the membrane in a horizontal orientation. To avoid this phenomenon, some solutions were proposed such as adding position restraints or rescaling protein-water van der Waals interactions [97]. In summary, Martini 3 is still in a testing phase by the scientific community. We are learning progressively how it behaves and how to interpret its outcomes by testing it on several different systems. In our case, we tested the rescaling procedure [97], but still observed TM1 ejections (data not shown).

In this context, how can we interpret the ejections of TM1 observed in our simulations with Martini 3? On one hand, this is unexpected since we are dealing with a TM helix. ^2^H NMR experiments have confirmed that a single TM helix containing a Lys in its center assumes a transmembrane topology regardless of the charge state [37]. On the other hand, we have to consider that there are 3 consecutive polar residues STK in the middle of TM1. Moreover, Lys 716 may be charged depending on the conditions. In the context of the full protein, which bears the HR1 cytoplasmic domain on its N-terminal side, these ejections are probably not realistic. But due to the simplified description of coarse-grained vs all-atom representation (fixed secondary structure, etc.), due to the absence of the cytoplasmic domains, this is how Martini 3 is warning us that something is going on with TM1: it is not an ideal helix for a hydrophobic environment like the center of a membrane. Yet, we demonstrated that TM1 destabilizes the membrane. If Lys716, were to be charged, this would be even more pronounced. This is interesting, because in the context of membrane fusion, a TM helix able to destabilize the membrane is clearly an asset, such as in SNARE proteins [98, 22]. We discuss this aspect further 2 sections below. Last, as stated by Sahoo *et al.*, we observed that the flanking residues matter, since the longer PEPT1-41 underwent less ejections than the shorter PEPT1-29 [35]. We also observed that charged Lys716 was ending in far more ejections than neutral Lys716. The ratio of ejections may thus be seen as a proxy towards the likeliness to perturb the membrane.

### Extensive multiscale simulations yield a new structural model of the Fzo1 TM domain improving over previous predictions

The TM domain model refined with our multiscale protocol shows significant deviations from the 2017 prediction [30]. The soluble part of the previous model could be validated by experimental mutation studies, while the TM1-TM2 contacts of the transmembrane part were based on an *ab initio* prediction by the PREDDIMER method [33] that could not be experimentally assessed at that time. Of the three PREDDIMER models created, the highest scoring conformation was selected, a right-handed antiparallel dimer characterized by a crossing angle of 119.7*^◦^*. In contrast, the model created here with Martini 3 is a left-handed antiparallel dimer with a crossing angle of −137.4*^◦^*, and the model refined with REMD has an angle of −161.4*^◦^*. In addition, the Martini 3 model shows involvement of the GX_3_G motif, while the refined model no longer shows interaction of these residues with TM1. This motif could thereby be free for other contacts such as homodimerization, or with another partner in the membrane. In addition, TM1 shows no matching interactions with the residues observed with PREDDIMER. T715 is most closely associated with the GX_3_G motif in the PREDDIMER model, whereas in the new model it is not T715 but L709, Y710, and G713, among others.

### An independent Fzo1-Ugo1 complex prediction matches the novel TM domain fold

The field of structure prediction has been revolutionized recently by the success of AlphaFold2 (AF2) [74, 75]. This observation prompted us to use AF2 to independently predict the structure of the TM domain of Fzo1.

Using AF2, we first predicted the TM domain of Fzo1 alone, then a whole Fzo1 monomer and a homodimer of Fzo1. All these predictions led to the two TM helices unstructured, with a low pLDDT (Predicted Local Distance Difference Test), which is the confidence value per residue (Supplementary Figure 13). This observations seems to echo the difficulty to predict membrane-inserted protein segments with AF2.

In yeast, Fzo1 has an important biological partner in the outer membrane named Ugo1 [99] which is involved in mitochondrial fusion [100, 101]. Interestingly, when two Fzo1 monomers and two Ugo1 monomers were subjected to AF2, we observed that the two TM helices were correctly predicted to interact with each other (Figure 5). Strikingly, compared to our best model from the REMD simulations (Figure 3D), the two structures are very close: the RMSD (on backbone C*α* atoms of TM1 and TM2 only) is as low as 0.27 nm, the promiscuous residues are very similar, and the crossing angles are close (−159*^◦^* for AF2, −163*^◦^* for our model) (Figure 5). Moreover, the AF2 model shows that the TM domain of each Fzo1 monomer interacts with each other around the TM helices (mainly TM2-TM2 contacts). This result is interesting because it suggests that Fzo1-Fzo1 dimerization may also involve contacts between the two TM domains. As the GX3G motif does not actually interact with TM1, these TM2 residues could be involved with the other monomer, even if we find closer interactions with other residues.

**Figure 5.**
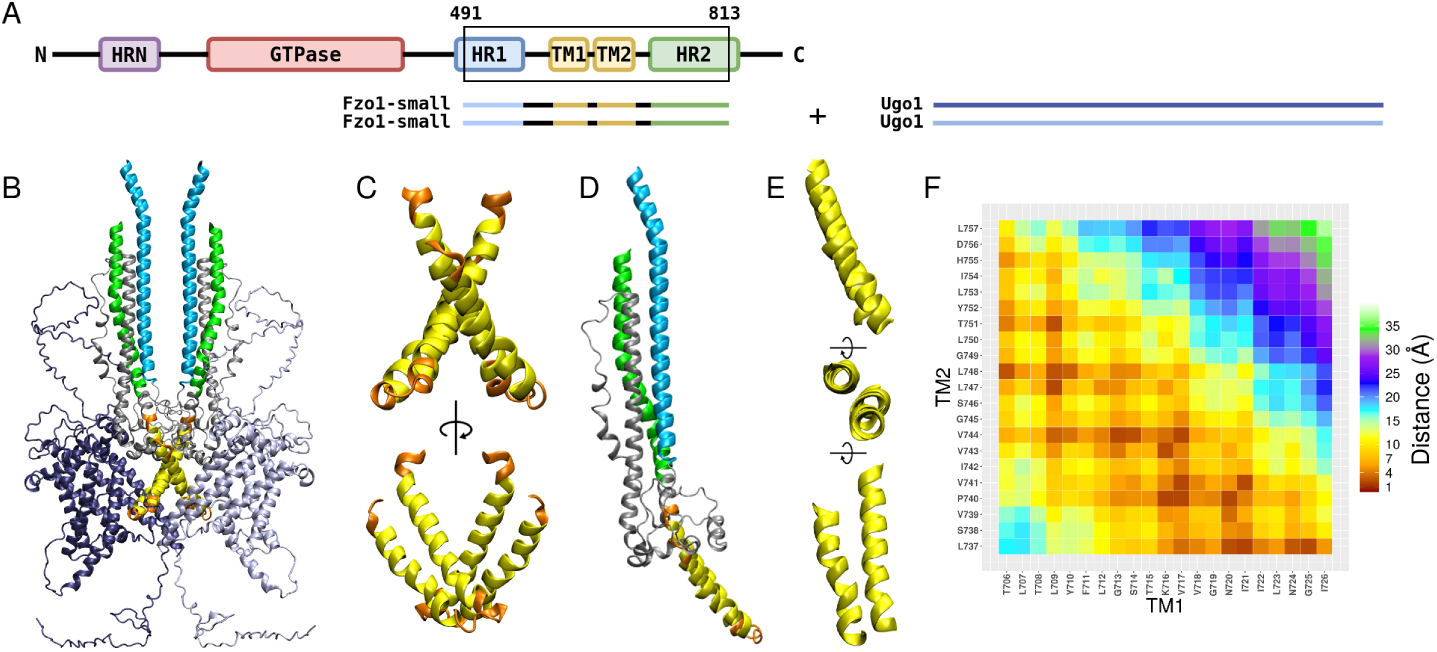
AlphaFold2 prediction of Fzo1 in interaction with Ugo1. (A) Scheme of the protocol. (B,C,D,E) The four proteins are in cartoon representation. In dark blue and light blue are the models of Ugo1. TM1 and TM2 are seen in yellow, the other residues used for the REMD are seen in orange, HR1 in blue and HR2 in green. The residues that do not belong to a specific domain are in light grey. (B) The overall result of AlphaFold. (C) Zoom on the TM domains of the Fzo1-small dimers. (D) One Fzo1-small monomer. (E) Zoom on TM1 and TM2 of Fzo1-small monomer. (F) Contact map of TM1 and TM2.

In summary, the overall agreement between our strategy and an artificial-intelligence-based method reinforces the pertinence of our physics-based model.

### Could the Fzo1 TM domain play an active role in membrane fusion?

The fusion process starts with the two membranes getting close to one another. The approach is followed by the formation of a stalk intermediate resulting from the mixing of the outer bilayer leaflets. Then a hemifused state follows prior to the final fused state. Outer leaflet mixing is possible when the hydrophobic areas are in contact with one another [102].

Recent experimental studies have shown the importance of lipid conformations in the initiation of the outer leaflets mixing. The ability of lipids to splay, i.e. expose a part of their hydrophobic tail towards the solvent also known as protrusion, was shown to trigger membrane fusion [103]. It was also shown that lipid packing defects, small hydrophobic areas exposed to the solvent, were qualitatively correlated to the nucleation rate of fusion [87]. Protrusions and packing defects are logically directly correlated on lipid composition and membrane curvature [104, 105, 62, 64]. These two parameters are thus critical in the initiation of membrane fusion.

Regarding membrane fusion mediated by proteins, it is now well established that their TM domains play a pivotal role [106]. In the well-studied case of SNARE proteins, the unique presence of a single TM helix in each leaflet catalyses outer leaflet mixing [98]. The TM helix sequence matters since some mutants induce less protrusions than the wild type [22]. In addition, the lipids are more perturbed near the TM helix than far away. Another well known example is the influenza fusion peptide [86]. Again, the TM sequence is decisive for the generation of protrusions around it (leading to membrane fusion), while some mutants are less efficient. A recent experimental study on model TM helices has also shown that the amount of lipid splay (i.e. protrusion) was directly correlated with membrane fusion [103]. In this example, membrane fusion also depended on the amino-acid sequence, a poly LV16 being more efficient than a poly L16 TM peptide. Similar to these examples, our results suggest a destabilization of the membrane promoted by the presence of the Fzo1 TM domain.

We discuss now the possible role of Lys716 in membrane fusion. First, this lysine is well conserved in fungi (Figure 13). In mammals, an arginine is rather found at this position. In organisms that have a full TM domain (not a paddle which only partially binds the membrane like in bacterial BDLP), it is interesting to note that evolution have conserved a basic residue in the middle of TM1, which raises questions about its role. Our experiments showed that mitochondrial respiration is disabled when Lys716 is mutated to an apolar residue (Leu or Val), but mutation to an Arg or Ser has no effect on respiration. Amino-acid polarity is thus required, but having a basic one is not mandatory. These data are in line with the membrane destabilization hypothesis, since a polar residue will anyway pertub the membrane (recall there are two other polar residues next to Lys716). However, these findings do not preclude the possibility of Lys716 positive charge to play a role. The experiments of Gleason *et al.* evaluated the pKa of a Lys in the middle of a TM helix to be around 6.2 at 50 *^◦^*C [37]. Correcting for temperature, the pKa was then estimated to be *∼* 6.5 at 37 *^◦^*C and *∼* 6.8 at 25 *^◦^*C. Given an acidic pH in yeast cytoplasm[107], these values would suggest that Lys716 could be, at least partially, protonated. The role of positively charged residues is not new in the context of membrane fusion. In the case of the SNARE fusion machinery, Lindau *et al.* examined the transmembrane domains in detail and hypothesized that well-placed charges could significantly assist membrane deformation and destructuring, thereby initiating the fusion process. In particular, they investigated the role of charged residues at the end of a Syb2 TM domain construct when pulled into the membrane [21]. It has further been suggested that charged motifs play an important role for fusion to occur, such as shown by the bilayer destabilization by a conserved membrane-embedded motif at the juxtamembrane region of the vesicle-anchored v-SNARE comprising several basic residues [108]. In TM1 of Fzo1, the Lys is in the middle of the TM helix, not flanking it, but it does not exclude that a similar mechanism may be operating. Overall, these observations echo the idea of a protonation-state induced switch of Lys716 to assist membrane fusion. However, the lack of effect when the Lys is mutated to a polar, but not titratable residue, is arguing against a direct effect on mitochondria fusion. Therefore, the ability of Lys716 to be protonated could be associated with the fusion of other membranes to which Fzo1 is associated. Besides mitochondrial fusion, we cannot exclude that this titration could be associated to another function.

## Conclusion

We have constructed an improved model of the TM domain of mitofusins using both coarse-grained and all-atom molecular dynamics. This model has been further confirmed using the deep-learning tool AlphaFold2. This model has revealed the role of Lys716 that is found in the middle of the membrane. The importance of this residue is confirmed by its evolutionary conservation and the effect of its mutations on mitochondrial fusion. Further studies will therefore be needed to explore the exact role of this residue in mitochondrial fusion but could also target other functions associated with mitofusins.

## Author Contributions

AT and PFJF contributed equally to this work. RV, AT and PFJF designed the research. RV carried out all simulations. RV, MB, AT and PFJF analyzed the simulation data. LC carried out the experiments. LC and MMC analysed the experiments. RV, MB, AT and PFJF wrote the article.

## Acknowledgments

This work was supported by Agence Nationale de la Recherche ANR-19-CE11-0018, MITOFUSION and by the “Initiative d’Excellence” program from the French State (Grant “DYNAMO”, ANR-11-LABX-0011). This work was granted access to the HPC resources of IDRIS under the allocation 2022-A0130701714 made by GENCI to MB. Computational work was performed using HPC resources from LBT-HPC thanks to support from Geoffrey Letessier. Drs Cathy Etchebest, Nicolas Floquet and Miguel Machuqueiro are acknowledged for stimulating discussions. Lucas Guémené is thanked for his help with experiments and Hanene Samia Ameur for testing Martini3 rescaling schemes.

## Supplementary Material

### Coarse-grained loop conformational sampling

#### System Building

The center of the first cluster obtained through Martini 3 simulations with a neutral Lys716 was selected for the following simulations. The lipids and solvent of the corresponding frame were also extracted. This dimer conformation was used as a starting point for the addition of the missing residues between PEPT1-41 and PEPT2 (containing respectively TM1 and TM2).

In order to add the missing residues between TM1 and TM2, a structure of the loop was built using Basic Builder 1.0.2 on the Mobyle server (https://mobyle.rpbs.univ-paris-diderot.fr) [45]. This structure was then converted to coarse-grained (CG) representation using the martinize2.py workflow module of the Martini 3 force field. As a first step, the residues at the end of the loop were used to align the structures of the dimer with the structure of the loop. Coordinates of residues Lys727 to Lys736 of the dimers were then removed and replaced by the coordinates of the structure available. Residues Arg695 to Thr701 included were also removed.

#### Simulation parameters

The GROMACS 2018.5 program was used to perform all MD simulations [39]. This system was then subjected to two energy minimization phase of 5000 and 10000 steps. Position restraints of 100000 *kJ.mol^−^*^1^.*nm^−^*^2^ were used for the TM1 and TM2 residues. Subsequently, all simulations were performed within the NPT ensemble. The velocity-rescaling thermostat [51] at 303.5K and the Berendsen barostat at 1bar [52] was applied for 500 ps of equilibration, with a time step of 0.01 ps. The position restraints for the TM1 and TM2 residues were lowered at 20000 *kJ.mol^−^*^1^.*nm^−^*^2^. A production run of 5 $*µ*$s followed at 303.5K using the velocity-rescaling thermostat [51] (lipid, water and proteins coupled separately) and at 1 bar using the Parrinello-Rahman barostat [53] (compressibility of 3.0 x 10*\^−^*^4^ bar). Pressure coupling was applied semi-isotropically. The position restraints for the same residues were fixed at 1000 *kJ.mol^−^*^1^.*nm^−^*^2^. A time step of 0.01 ps was used with the leapfrog integrator. Lennard-Jones interactions were cutoff at 1.1 nm. Bond lengths were constrained using the LINCS algorithm [54]. The reaction-field method [55] was used for evaluating electrostatic interactions, with a Coulomb distance cutoff of 1.1 nm, a relative dielectric constant of 15. The neighbor list was updated every 20 steps.

At the end of this stage, we thus have a coarse-grained system with TM1 and TM2 connected by the loop and embedded in the bilayer surounded by solvent and ions.

#### Clustering

The loop was submitted to a conformation based clustering. The first 200 ns of the production were ignored, resulting in a total of 9601 frames. The frames underwent a clustering with the GROMOS method [56] using a cutoff of 0.2 nm (in order to obtain around 5 clusters representing 80-90 % of all conformations). The center conformation of the first cluster was then selected for the next step.

## Supplementary Results

**Figure 6.**
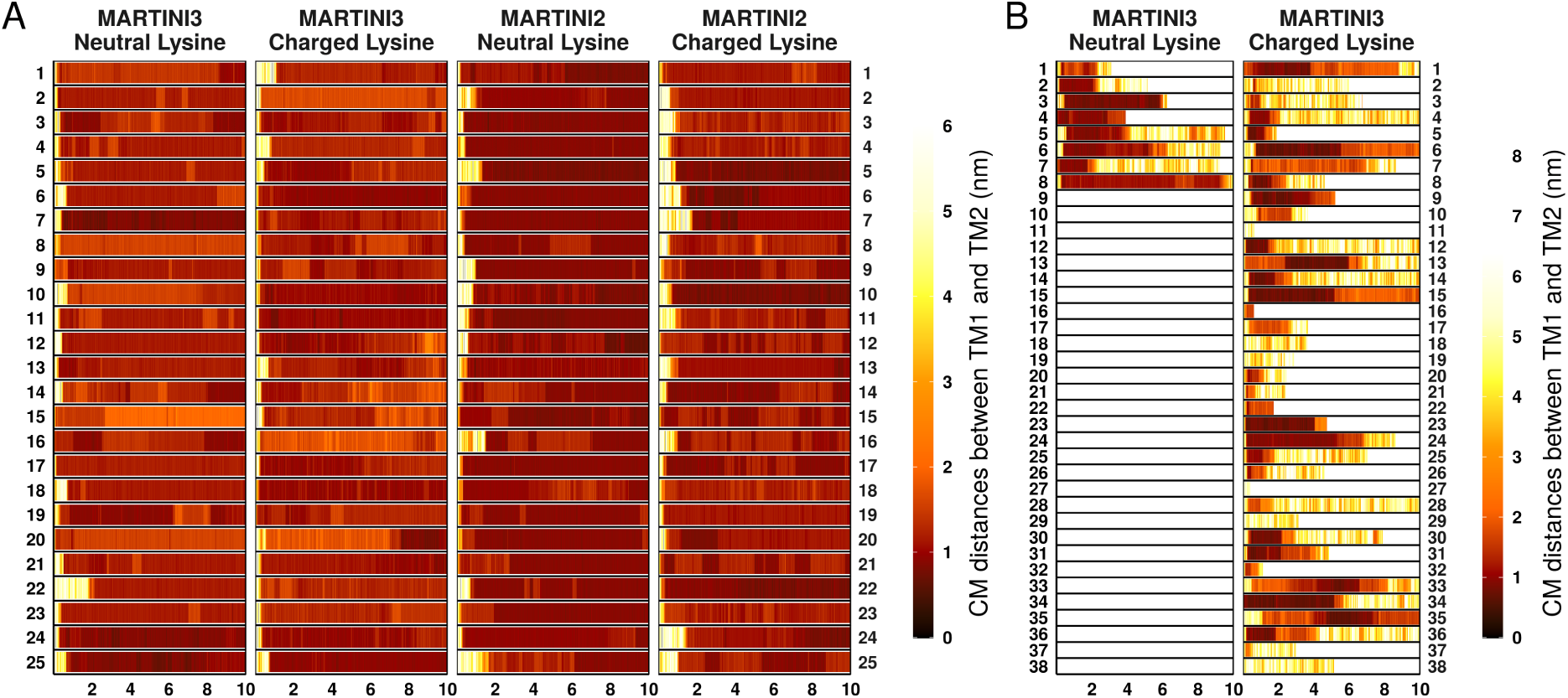
Distance between the center of mass of the two TM domains, as a function of time. (A) Results for Martini 2 and Martini 3 systems. (B) Results for Martini 3 simulations, where TM1 exiting the membrane was observed.

**Figure 7.**
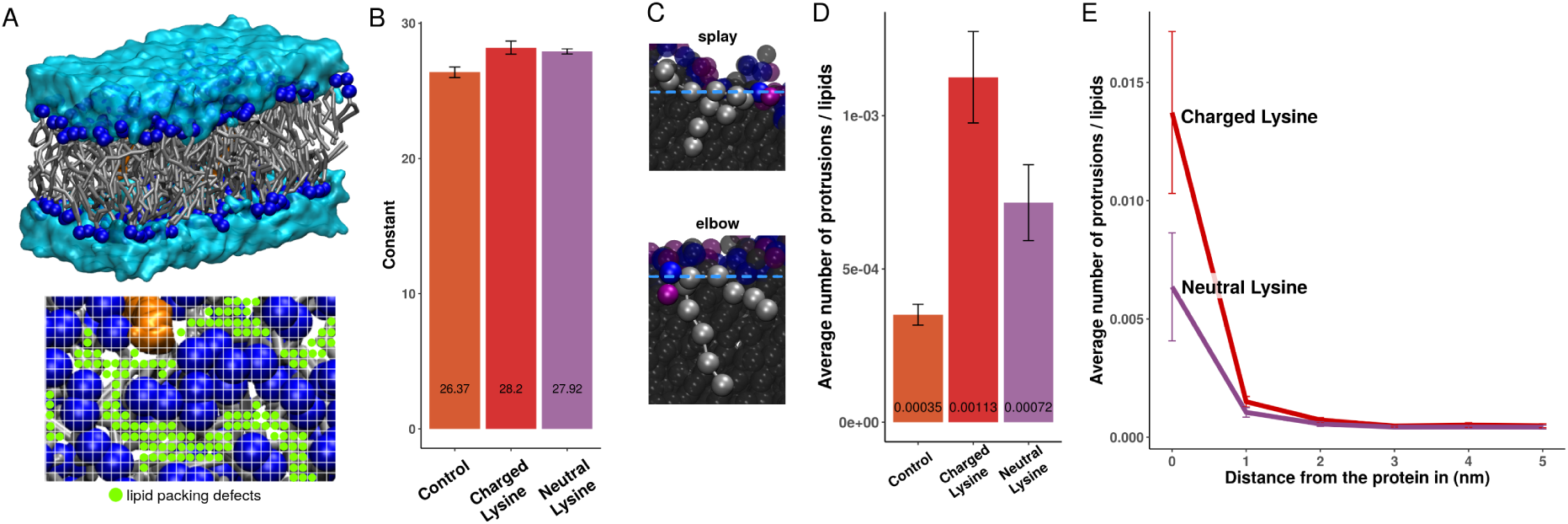
Caption

**Figure 8.**
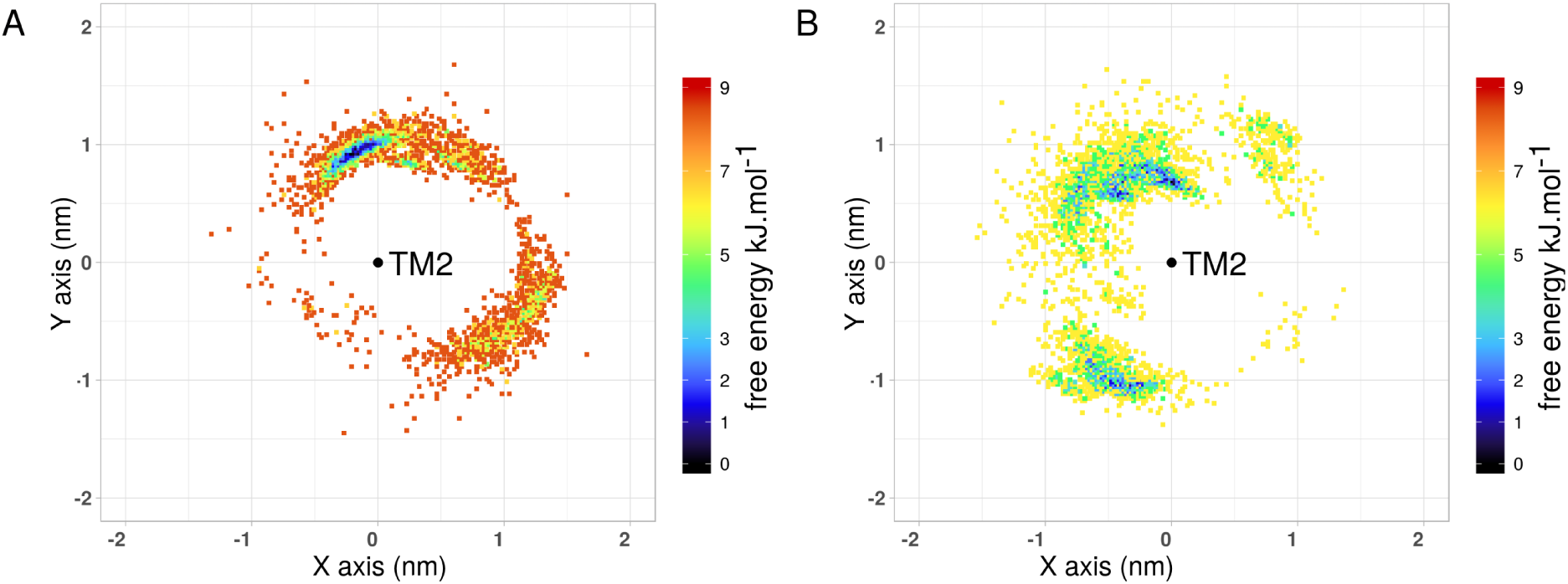
Position of TM1 relative to TM2 colored as a function of the free energy. (A) Lys716 charged. (B) Lys716 neutral.

**Figure 9.**
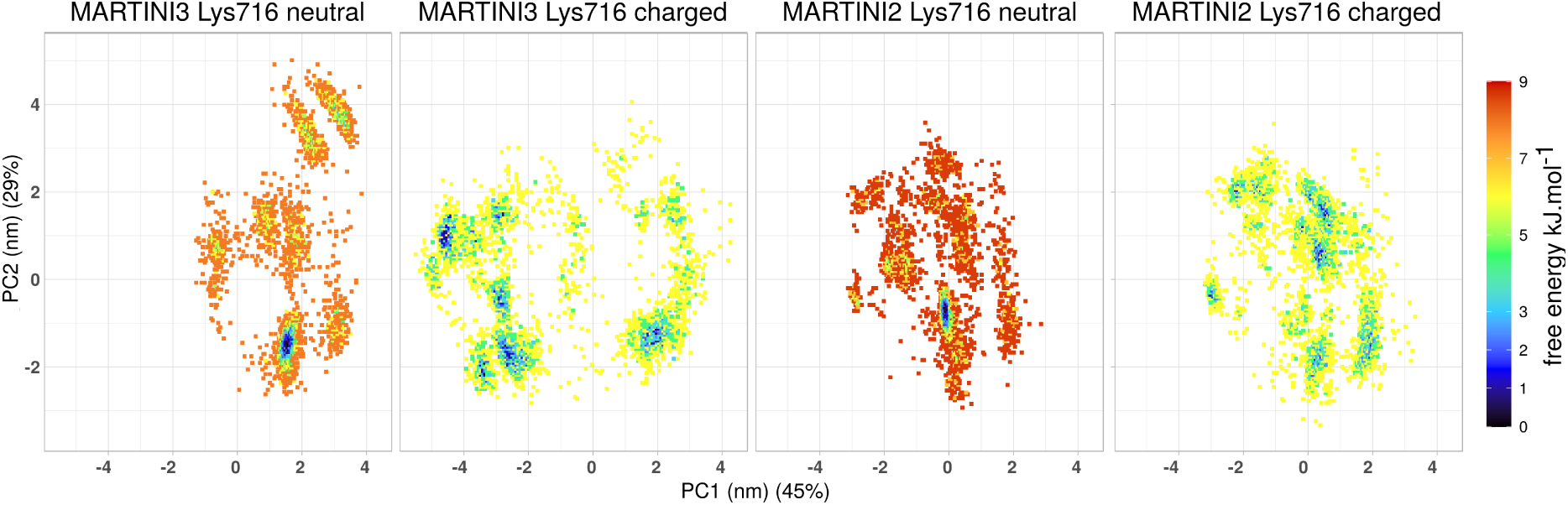
PCA colored as a function of the free energy.

**Figure 10.**
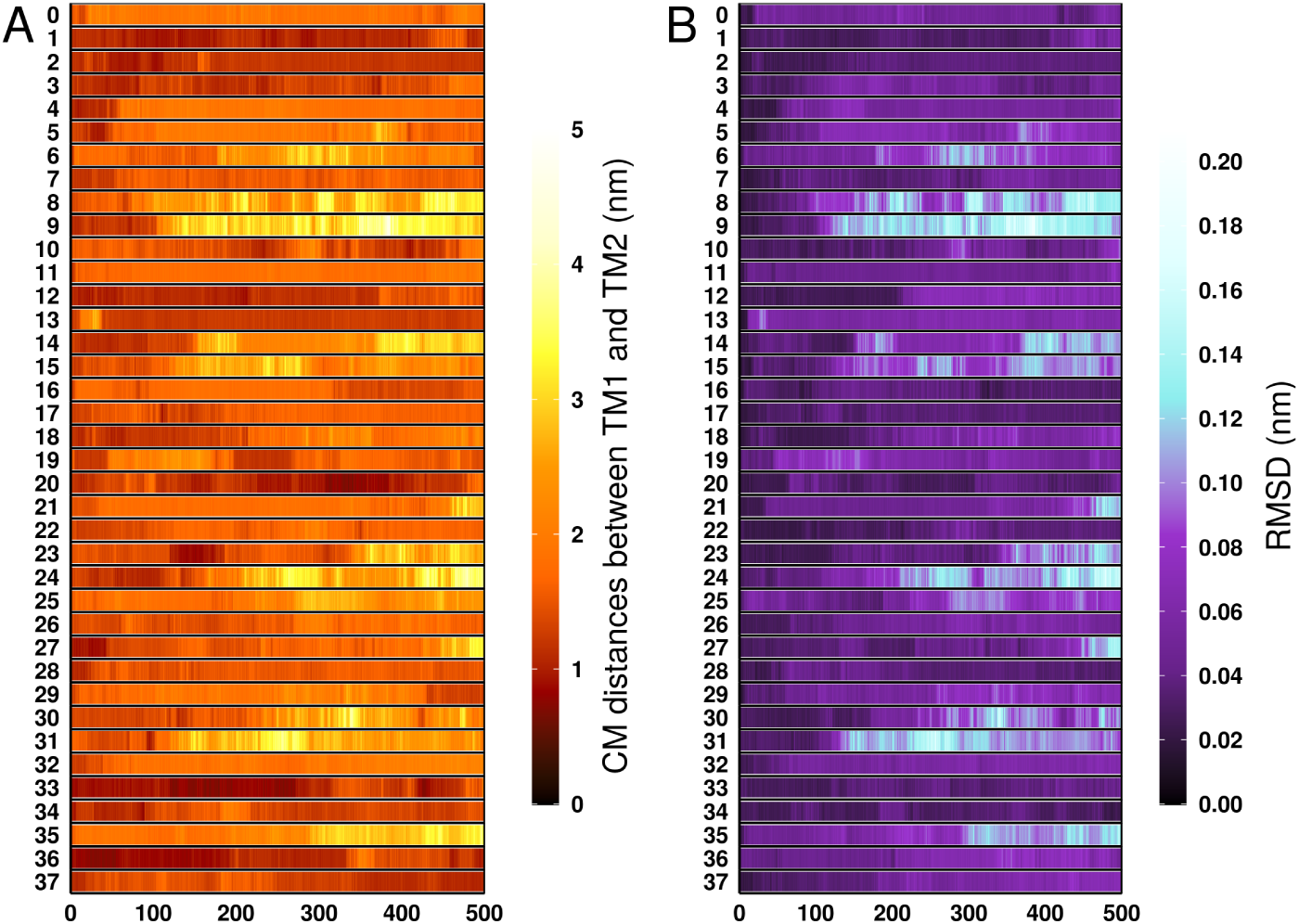
Supplementary results of the REMD. (A) Distance between the center of mass of TM1 and TM2 as a function of time for for all replicas. (B) RMSD of TM1 and TM2 (considering C*_α_* only) with respect to the starting structure as a function of time.

**Figure 11.**
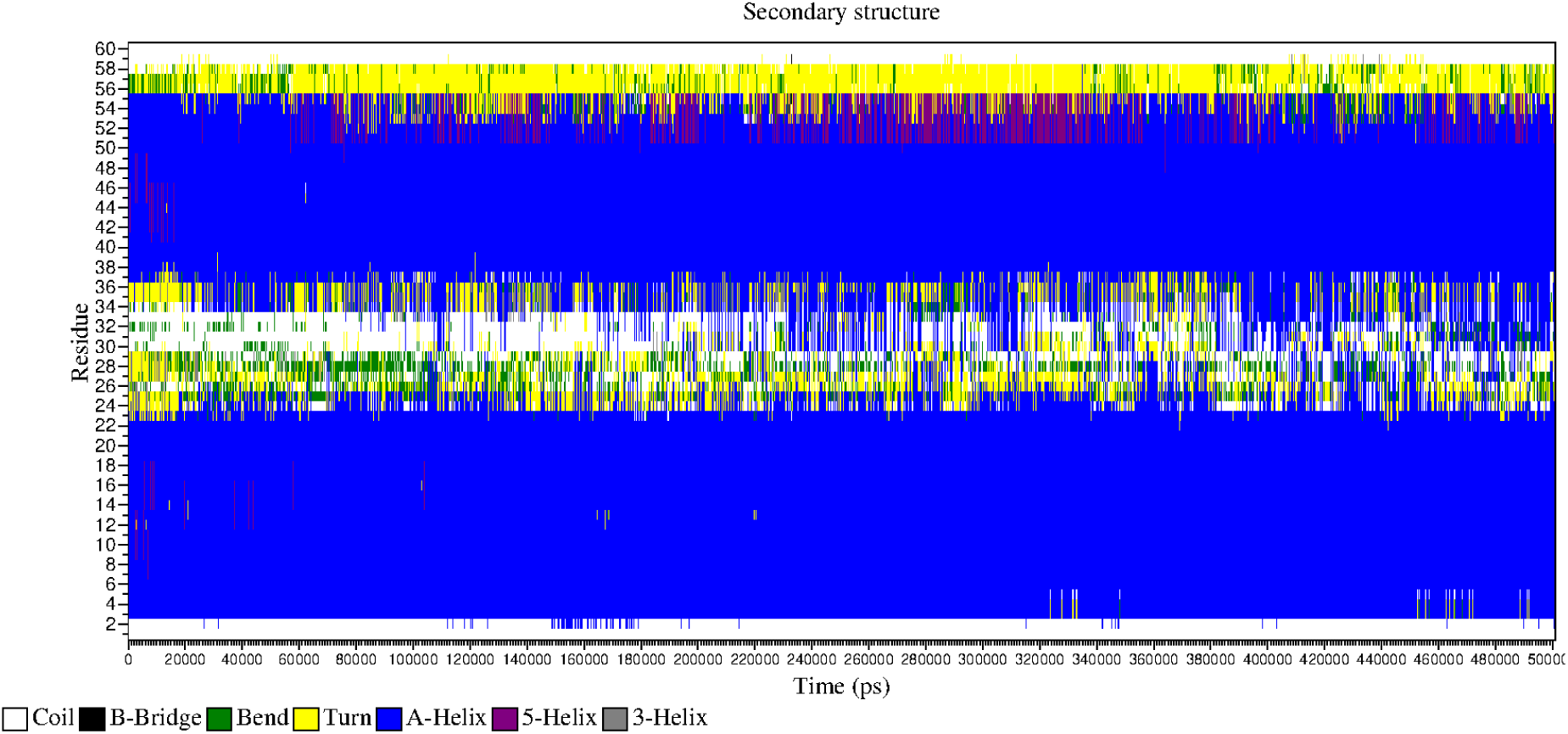
Supplementary results of the REMD. Secondary structure of the ensemble of structures at 303.15 K.

**Figure 12.**
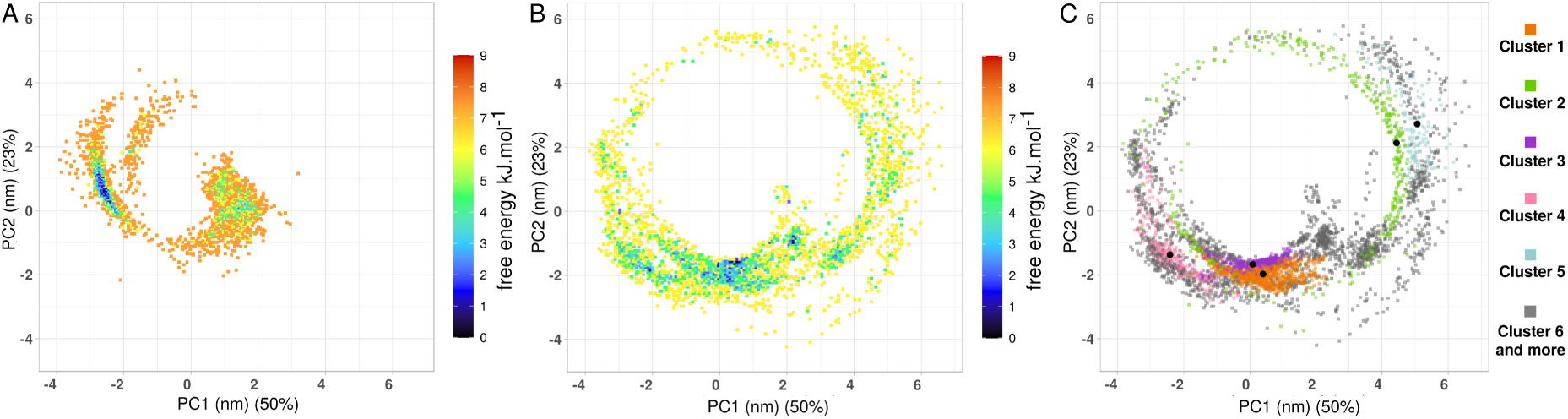
PCA results of the REMD. (A) Martini 3 Lys716 neutral structures colored as function of free energy. (B) REMD structures colored as function of free energy. (C) REMD structures colored as a function of the cluster number to which they belong to.

**Figure 13.**
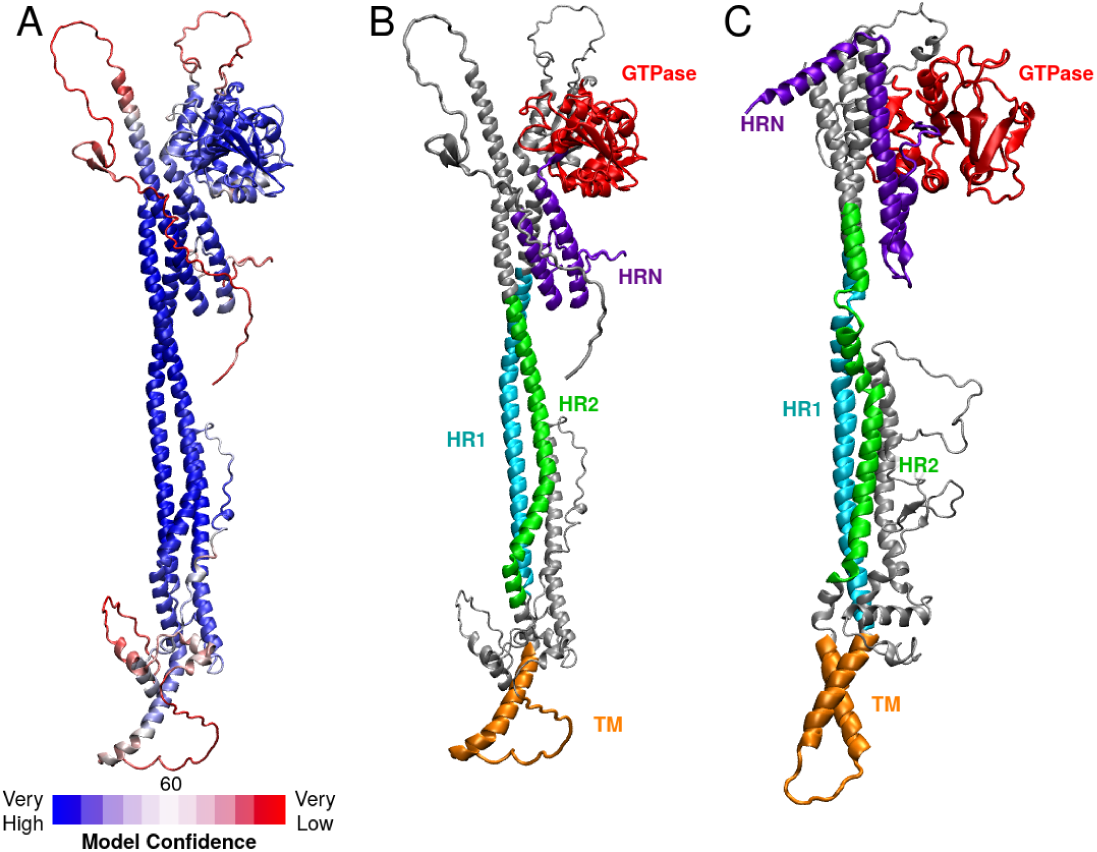
Fzo1 model of Alphafold. The model is shown in cartoon representation. (A) Colored as a function of plddt. (B) The domains are colored, and the residues outside of the domains are in grey. (C) Model built in 2017 [30], for comparison purposes.

**Figure 14.**
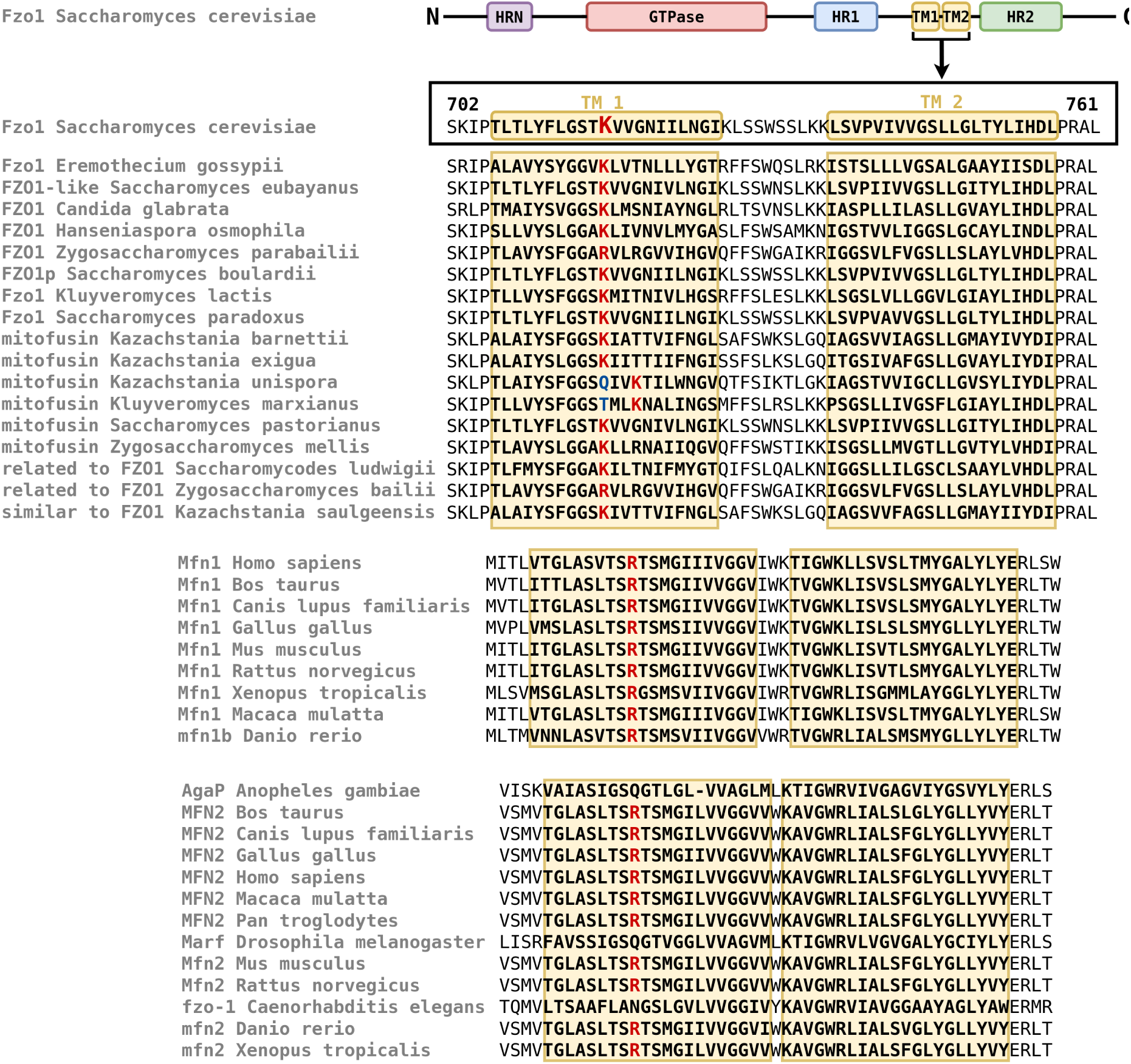
**Mitofusins TM sequences alignments.**

## References

1. Benedikt Westermann. Mitochondrial fusion and fission in cell life and death. Nat. Rev. Mol. Cell Biol., 11(12):872–884, 2010.

2. Rajesh Ramachandran. Mitochondrial dynamics: The dynamin superfamily and execution by collusion. Semin. Cell Dev. Biol., 76:201–212, 2018.

3. Naotada Ishihara, Yuka Eura, and Katsuyoshi Mihara. Mitofusin 1 and 2 play distinct roles in mitochondrial fusion reactions via GTPase activity. Journal of Cell Science, 117(26):6535–6546, 2004.

4. Manuel Rojo, Frédéric Legros, Danielle Chateau, and Anne Lombés. Membrane topology and mitochondrial targeting of mitofusins, ubiquitous mammalian homologs of the transmembrane GTPase fzo. J. Cell. Sci., 115:1663–1674, 2002.

5. G. J. Hermann, J. W. Thatcher, J. P. Mills, K. G. Hales, M. T. Fuller, J. Nunnari, and J. M. Shaw. Mitochondrial fusion in yeast requires the transmembrane GTPase fzo1p. J. Cell Biol., 143(2):359–373, 1998.

6. Yu-Lu Cao, Shuxia Meng, Yang Chen, Jian-Xiong Feng, Dong-Dong Gu, Bing Yu, Yu-Jie Li, Jin-Yu Yang, Shuang Liao, David C. Chan, and Song Gao. Mfn1 structures reveal nucleotide-triggered dimerization critical for mitochondrial fusion. Nature, 542:372–376, 2017.

7. Liming Yan, Yuanbo Qi, Xiaofang Huang, Caiting Yu, Lan Lan, Xiangyang Guo, Zihe Rao, Junjie Hu, and Zhiyong Lou. Structural basis for gtp hydrolysis and conformational change of mfn1 in mediated membrane fusion. Nat. Struct. Mol., 25:233–243, 2017.

8. E. D. Wong, J. A. Wagner, S. W. Gorsich, J. M. McCaffery, J. M. Shaw, and J. Nunnari. The dynamin-related GTPase, mgm1p, is an intermembrane space protein required for maintenance of fusion competent mitochondria. J. Cell Biol., 151(2):341–352, 2000.

9. Christian Frezza, Sara Cipolat, Olga Martins de Brito, Massimo Micaroni, Galina V. Bez-noussenko, Tomasz Rudka, Davide Bartoli, Roman S. Polishuck, Nika N. Danial, Bart De Strooper, and Luca Scorrano. OPA1 controls apoptotic cristae remodeling independently from mitochondrial fusion. Cell, 126(1):177–189, 2006.

10. Hsiuchen Chen and David C. Chan. Mitochondrial dynamics–fusion, fission, movement, and mitophagy–in neurodegenerative diseases. Hum Mol Genet, 18:R169–R176, 2009.

11. Andrew B. Knott, Guy Perkins, Robert Schwarzenbacher, and Ella Bossy-Wetzel. Mitochondrial fragmentation in neurodegeneration. Nature Reviews Neuroscience, 9(7):505–518, 2008.

12. Keshav K Singh and Josephine S. Modica-Napolitano. Special issue: Mitochondria in cancer. Seminars in Cancer Biology, 47:iv–vi, 2017. Mitochondria in Cancer.

13. Alessandro Allegra, Vanessa Innao, Andrea Gaetano Allegra, and Caterina Musolino. Relationship between mitofusin 2 and cancer. Advances in Protein Chemistry and Structural Biology, 116:209–236, 2019.

14. Fatemeh Darvish Moghaddam, Peyman Mortazavi, Saeed Hamedi, Mohammad Nabiuni, and Nastaran Hassanzadeh Roodbari. Apoptotic effects of melittin on 4t1 breast cancer cell line is associated with up regulation of mfn1 and drp1 mrna expression. Anticancer Agents Med Chem, 20(7):790–799, 2020.

15. Zhong Zhang, Ting-E Li, Min Chen, and, et al. Mfn1-dependent alteration of mitochondrial dynamics drives hepatocellular carcinoma metastasis by glucose metabolic reprogramming. British Journal of Cancer, 122:209–220, feb 2020.

16. Rehman Jalees, Zhang Hannah J, Toth Peter T, Zhang Yanmin, Marsboom Glenn, Hong Zhigang, Salgia Ravi, Husain Aliya N, Wietholt Christian, and Archer Stephen L. Inhibition of mitochondrial fission prevents cell cycle progression in lung cancer. FASEB J., 26(5):0892–6638, 2012. PMID: 22321727.

17. Ahn Sung Yong, Song Jiwon, Kim Yu Cheon, Kim Myoung Hee, and Hyun Young-Min. Mitofusin-2 promotes the epithelial-mesenchymal transition-induced cervical cancer progression. Immune Netw, 21, 2021.

18. Xiaofei Cheng, Yanqing Li, and Fanlong Liu. Prognostic impact of mitofusin 2 expression in colon cancer. Translational Cancer Research, 11(10), 2022.

19. S.M.E. Feely, M. Laura, C.E. Siskind, S. Sottile, M. Davis, V.S. Gibbons, M.M. Reilly, and M.E. Shy. Mfn2 mutations cause severe phenotypes in most patients with cmt2a. Neurology, 76(20):1690–1696, 2011.

20. Stephan Z”uchner, Irina V Mersiyanova, Maria Muglia, Neda Bissar-Tadmouri, Jennifer Rochelle, Ece L Dadali, Mario Zappia, Eva Nelis, Antonella Patitucci, Jan Senderek, et al. Mutations in the mitochondrial gtpase mitofusin 2 cause charcot-marie-tooth neuropathy type 2a. Nat Genet, 36(5):449–451, 2004. Erratum in: Nat Genet. 2004 Jun;36(6):660. Battologlu E [corrected to Battaloglu E].

21. Manfred Lindau, Benjamin A. Hall, Alan Chetwynd, Oliver Beckstein, and Mark S. P. Sansom. Coarse-Grain Simulations Reveal Movement of the Synaptobrevin C-Terminus in Response to Piconewton Forces. Biophysical Journal, 103(5):959–969, September 2012.

22. Jing Han, Kristyna Pluhackova, Dieter Bruns, and Rainer A. Böckmann. Synaptobrevin transmembrane domain determines the structure and dynamics of the snare motif and the linker region. Biochimica et Biophysica Acta (BBA) - Biomembranes, 1858(4):855–865, 2016.

23. Jan-Dirk Wehland, Antonina S. Lygina, Pawan Kumar, Samit Guha, Barbara E. Hubrich, Reinhard Jahn, and Ulf Diederichsen. Role of the transmembrane domain in snare protein mediated membrane fusion: peptide nucleic acid/peptide model systems. Mol. BioSyst., 12:2770–2776, 2016.

24. Madhurima Dhara, Antonio Yarzagaray, Mazen Makke, Barbara Schindeldecker, Yvonne Schwarz, Ahmed Shaaban, Satyan Sharma, Rainer A Böckmann, Manfred Lindau, Ralf Mohrmann, and Dieter Bruns. v-snare transmembrane domains function as catalysts for vesicle fusion. eLife, 5:e17571, jun 2016.

25. Greg J. Hermann, John W. Thatcher, John P. Mills, Karen G. Hales, Margaret T. Fuller, Jodi Nunnari, and Janet M. Shaw. Mitochondrial fusion in yeast requires the transmembrane gtpase fzo1p. J. Cell Biol., 143:359–373, 1998.

26. Erik E. Griffin and David C. Chan. Domain interactions within fzo1 oligomers are essential for mitochondrial fusion. J. Biol. Chem., 281(24):16599–16606, 2006.

27. Harry H. Low and Jan Löwe. A bacterial dynamin-like protein. Nature, 444:766–769, 2006.

28. Harry H. Low, Carsten Sachse, Linda A. Amos, and Jan Löwe. Structure of a bacterial dynamin-like protein lipid tube provides a mechanism for assembly and membrane curving. J. Cell Biol., 139:1342–1352, 2009.

29. Stefan Fritz, Doron Rapaport, Elisabeth Klanner, Walter Neupert, and Benedikt Westermann. Connection of the mitochondrial outer and inner membranes by fzo1 is critical for organellar fusion. J. Cell Bio., 152:683–692, 2001.

30. Dario De Vecchis, Laetitia Cavellini, Marc Baaden, Jér^ome Hénin, Mickaël M. Cohen, and Antoine Taly. A membrane-inserted structural model of the yeast mitofusin fzo1. Scientific Reports, 7(1):10217, 2017.

31. Astrid Brandner, Dario De Vecchis, Marc Baaden, Mickael M. Cohen, and Antoine Taly. Physics-based oligomeric models of the yeast mitofusin fzo1 at the molecular scale in the context of membrane docking. Mitochondrion, 49:234–244, 2019.

32. Dario De Vecchis, Astrid Brandner, Marc Baaden, Mickael M. Cohen, and Antoine Taly. A molecular perspective on mitochondrial membrane fusion: From the key players to oligomerization and tethering of mitofusin. J. Membr. Biol., 252(4):293–306, 2019.

33. Anton A. Polyansky, Anton O. Chugunov, Pavel E. Volynsky, Nikolay A. Krylov, Dmitry E. Nolde, and Roman G. Efremov. Preddimer: a web server for prediction of transmemrane helical dimers. Bioinformatics, 30:889–890, 2014.

34. Paulo C. T. Souza, Roberto Alessandri, Julien Barnoud, and, et al. Martini 3: a general purpose force field for coarse-grained molecular dynamics. Nat Methods, 18:382–388, 2021.

35. Amita R Sahoo, Paulo CT Souza, Zhiyuan Meng, and Matthias Buck. Transmembrane dimers of type 1 receptors sample alternate configurations: Md simulations using coarse grain martini 3 versus alphafold2 multimer. Structure, 31(6):735–745, 2023.

36. Justin L. MacCallum, W. F. Drew Bennett, and D. Peter Tieleman. Distribution of amino acids in a lipid bilayer from computer simulations. Biophysical Journal, 94(9):3393–3404, 2008.

37. Nicholas J. Gleason, Vitaly V. Vostrikov, Denise V. Greathouse, and Roger E. Koeppe. Buried lysine, but not arginine, titrates and alters transmembrane helix tilt. Proceedings of the National Academy of Sciences, 110(5):1692–1695, 2013.

38. Afra Panahi and Charles L. III Brooks. Membrane environment modulates the pka values of transmembrane helices. The Journal of Physical Chemistry B, 119(13):4601–4607, 2015. PMID: 25734901.

39. Mark James Abraham, Teemu Murtola, Roland Schulz, Szilárd Páll, Jeremy C Smith, Berk Hess, and Erik Lindahl. Gromacs: High performance molecular simulations through multi-level parallelism from laptops to supercomputers. SoftwareX, 1:19–25, 2015.

40. UniProt Consortium. UniProt: a worldwide hub of protein knowledge. Nucleic acids research, 47(D1):D506–D515, 2018.

41. Daniel WA Buchan and David T Jones. The PSIPRED protein analysis workbench: 20 years on. Nucleic acids research, 47(W1):W402–W407, 2019.

42. David T Jones. Protein secondary structure prediction based on position-specific scoring matrices. Journal of molecular biology, 292(2):195–202, 1999.

43. Anders Krogh, BjoÈrn Larsson, Gunnar Von Heijne, and Erik LL Sonnhammer. Predicting transmembrane protein topology with a hidden Markov model: application to complete genomes. Journal of molecular biology, 305(3):567–580, 2001.

44. E. L. Sonnhammer, G. von Heijne, and A. Krogh. A hidden markov model for predicting transmembrane helices in protein sequences. Proc Int Conf Intell Syst Mol Biol, 6:175–182.

45. Bertrand Néron, Hervé Ménager, Corinne Maufrais, Nicolas Joly, Julien Maupetit, Sébastien Letort, Sébastien ans Carrere, Pierre Tuffery, and Catherine Letondal. Mobyle: a new full web bioinformatics framework. 25(22):3005–3011.

46. Sunhwan Jo, Taehoon Kim, Vidyashankara G. Iyer, and Wonpil Im. Charmm-gui: A web-based graphical user interface for charmm. Journal of Computational Chemistry, 29:1859–1865, 2008.

47. Yifei Qi, Helgi I. Inǵolfsson, Xi Cheng, Jumin Lee, Siewert J. Marrink, and Wonpil Im. Charmmgui martini maker for coarse-grained simulations with the martini force field. J. Chem. Theory and Comput., 11(9):4486–4494, 2015.

48. Djurre H. de Jong, Gurpreet Singh, W. F. Drew Bennett, Clement Arnarez, Tsjerk A. Wassenaar, Lars V. Schäfer, Xavier Periole, D. Peter Tieleman, and Siewert J. Marrink. Improved parameters for the martini coarse-grained protein force field. Journal of chemical theory and computation, 9(1):687–697, 2012.

49. Siewert J Marrink, H Jelger Risselada, Serge Yefimov, D Peter Tieleman, and Alex H De Vries. The martini force field: coarse grained model for biomolecular simulations. The journal of physical chemistry B, 111(27):7812–7824, 2007.

50. Luca Monticelli, Senthil K Kandasamy, Xavier Periole, Ronald G Larson, D Peter Tieleman, and Siewert-Jan Marrink. The martini coarse-grained force field: extension to proteins. Journal of chemical theory and computation, 4(5):819–834, 2008.

51. Giovanni Bussia, Davide Donadio, and Michele Parrinello. Canonical sampling through velocity rescaling. J. Chem. Phys., 126:014101–014107, 2007.

52. H. J. C. Berendsen, J. P. M. Postma, W. F. van Gunsteren, A. DiNola, and J. R. Haak. Molecular dynamics with coupling to an external bath. J. Chem. Phys., 81:3684–3690, 1984.

53. M. Parrinello and A. Rahman. Polymorphic transitions in single crystals: A new molecular dynamics method. J. Appl. Phys., 52:7182–7190, 1981.

54. Berk Hess, Henk Bekker, Herman J. C. Berendsen, and Johannes G. E. M. Fraaije. LINCS: A linear constraint solver for molecular simulations. Journal of Computational Chemistry, 18(12):1463–1472, 1997.

55. Ilario G. Tironi, René Sperb, Paul E. Smith, and Wilfred F. van Gunsteren. A generalized reaction field method for molecular dynamics simulations. J. Chem. Phys., 102(13):5451–5459, 1995.

56. Xavier Daura, Karl Gademann, Bernhard Jaun, Wilfred F. Seebach, Dieter ans van Gunsteren, and Alan E. Mark. Peptide folding: When simulation meets experiment. Angew. Chem. Int. Ed., 38:236–240, 1999.

57. Cyrus Chothia, Michael Levitt, and Douglas Richardson. Helix to helix packing in proteins. 145:215–250.

58. Naveen Michaud-Agrawal, Elizabeth J. Denning, Thomas B. Woolf, and Oliver Beckstein. MDAnalysis: A toolkit for the analysis of molecular dynamics simulations. 32(10):2319–2327.

59. Richard J. Gowers, Max Linke, Jonathan Barnoud, Tyler J. E. Reddy, Manuel N. Melo, Sean L. Seyler, Jan Domański, David L. Dotson, Sébastien Buchoux, Ian M. Kenney, and Oliver Beckstein. MDAnalysis: A python package for the rapid analysis of molecular dynamics simulations. pages 98–105.

60. Hadley Wickham. ggplot2: Elegant Graphics for Data Analysis. Springer-Verlag New York, 2016.

61. Lydie Vamparys, Romain Gautier, Stefano Vanni, W F Drew Bennett, D Peter Tieleman, Bruno Antonny, Catherine Etchebest, and Patrick F J Fuchs. Conical lipids in flat bilayers induce packing defects similar to that induced by positive curvature. Biophysical journal, 104(3):585—593, February 2013.

62. Romain Gautier, Amélie Bacle, Marion L. Tiberti, Patrick F. Fuchs, Stefano Vanni, and Bruno Antonny. Packmem: A versatile tool to compute and visualize interfacial packing defects in lipid bilayers. Biophysical Journal, 115(3):436–444, 2018.

63. Per Larsson and Peter M. Kasson. Lipid tail protrusion in simulations predicts fusogenic activity of influenza fusion peptide mutants and conformational models. PLoS Comput Biol, 9(3):e1002950, 2013.

64. Mukarram A.Tahir, Reid C.Van Lehn, S.H. Choi, and Alfredo Alexander-Katz. Solvent-exposed lipid tail protrusions depend on lipid membrane composition and curvature. Biochimica et Biophysica Acta (BBA) -Biomembranes, 1858(6):1207–1215, 2016.

65. Jing Huang, Sarah Rauscher, Grzegorz Nawrocki, Ting Ran, Michael Feig, Bert L. de Groot, Helmut Grubmüller, and Alexander D. MacKerell. CHARMM36m: An improved force field for folded and intrinsically disordered proteins. 14(1):71–73.

66. Jeffery B. Klauda, Richard M. Venable, J. Alfredo Freites, Joseph W. O’Connor, Douglas J. Tobias, Carlos Mondragon-Ramirez, Igor Vorobyov, Alexander D. Jr. MacKerell, and Richard W. Pastor. Update of the charmm all-atom additive force field for lipids: Validation on six lipid types. The Journal of Physical Chemistry B, 114(23):7830–7843, 2010. PMID: 20496934.

67. Tsjerk A. Wassenaar, Kristyna Pluhackova, Rainer A. Böckmann, Siewert J. Marrink, and D. Peter Tieleman. Going backward: A flexible geometric approach to reverse transformation from coarse grained to atomistic models. Journal of Chemical Theory and Computation, 10(2):676–690, 2014. Relation: https://www.rug.nl/ Rights: University of Groningen, Groningen Biomolecular Sciences and Biotechnology Institute.

68. Tom Darden, Darrin York, and Lee Pedersen. Particle mesh ewald: An n log (n) method for ewald sums in large systems. The Journal of chemical physics, 98(12):10089–10092, 1993.

69. Ulrich Essmann, Lalith Perera, Max L Berkowitz, Tom Darden, Hsing Lee, and Lee G Pedersen. A smooth particle mesh ewald method. The Journal of chemical physics, 103(19):8577–8593, 1995.

70. Berk Hess, Henk Bekker, Herman JC Berendsen, and Johannes GEM Fraaije. Lincs: A linear constraint solver for molecular simulations. Journal of computational chemistry, 18(12):1463– 1472, 1997.

71. Shuichi Miyamoto and Peter A Kollman. Settle: An analytical version of the shake and rattle algorithm for rigid water models. Journal of computational chemistry, 13(8):952–962, 1992.

72. Yuji Sugita and Yuko Okamoto. Replica-exchange molecular dynamics method for protein folding. Chemical physics letters, 314(1-2):141–151, 1999.

73. Alexandra Patriksson and David van der Spoel. A temperature predictor for parallel tempering simulations. Physical Chemistry Chemical Physics, 10(15):2073–2077, 2008.

74. John Jumper, Richard Evans, Alexander Pritzel, et al. Highly accurate protein structure prediction with alphafold. Nature, 596:583–589, 2021.

75. Richard Evans and et al. Protein complex prediction with alphafold-multimer. biorxiv, 2021.

76. Milot Mirdita, Konstantin Schütze, Yuki Moriwaki, Lim Heo Heo, Sergey Ovchinnikov, and Martin Steinegger. Colabfold: Making protein folding accessible to all. Nature Methods, 2022.

77. F Sherman, G Fink, and J Hicks. Methods in yeast genetics: a laboratory course manual, 1987.

78. Christiane Volland, Daniele Urban-Grimal, Gerard Geraud, and Rosine Haguenauer-Tsapis. Endocytosis and degradation of the yeast uracil permease under adverse conditions. Journal of Biological Chemistry, 269(13):9833–9841, 1994.

79. Robert S Sikorski and Philip Hieter. A system of shuttle vectors and yeast host strains designed for efficient manipulation of dna in saccharomyces cerevisiae. Genetics, 122(1):19–27, 1989.

80. Mickael M Cohen, Elizabeth A Amiott, Adam R Day, Guillaume P Leboucher, Erin N Pryce, Michael H Glickman, J Michael McCaffery, Janet M Shaw, and Allan M Weissman. Sequential requirements for the gtpase domain of the mitofusin fzo1 and the ubiquitin ligase scfmdm30 in mitochondrial outer membrane fusion. Journal of cell science, 124(9):1403–1410, 2011.

81. Bettina Brosig and Dieter Langosch. The dimerization motif of the glycophorin a transmembrane segment in membrane: importance of glycine residues. 7:1052–1056.

82. Pierre Hubert, Paul Sawma, Jean-Pierre Duneau, Jonathan Khao, Jéler^ome Dominique Hénin, Bagnard, and James Sturgis. Single-spanning transmembrane domains in cell growth and cell-cell interactions: More than meets the eye? Cell Adh Migr, 4:313–324.

83. R. F. S. Walters and W. F. DeGrado. Helix packing motifs in membrane proteins. Proc. Natl. Acad., 103:13658–13663.

84. Shao-Qing Zhang, Daniel W. Kulp, Chaim A. Schramm, Marco Mravic, Ilan Samish, and William F. DeGrado. The membrane- and soluble-protein helix-helix interactome: similar geometry via different interactions. Structure, 23:527–541.

85. Frédéric Pincet, Luc Lebeau, and Sophie Cribier. Short-range specific forces are able to induce hemifusion. European Biophysics Journal, 30:91–97, 2001.

86. Per Larsson and Peter M Kasson. Lipid tail protrusion in simulations predicts fusogenic activity of influenza fusion peptide mutants and conformational models. PLoS Computational Biology, 9(3):e1002950, 13.

87. Claire Fraņcois-Martin, Amélie Bacle, James E. Rothman, Patrick F. J. Fuchs, and Frédéric Pincet. Cooperation of conical and polyunsaturated lipids to regulate initiation and processing of membrane fusion. Frontiers in Molecular Biosciences, 8, 2021.

88. Filip Rolland, Joris Winderickx, and Johan M Thevelein. Glucose-sensing and-signalling mechanisms in yeast. FEMS yeast research, 2(2):183–201, 2002.

89. Juana M Gancedo. Yeast carbon catabolite repression. Microbiology and molecular biology reviews, 62(2):334–361, 1998.

90. Greg J Hermann, John W Thatcher, John P Mills, Karen G Hales, Margaret T Fuller, Jodi Nunnari, and Janet M Shaw. Mitochondrial fusion in yeast requires the transmembrane gtpase fzo1p. The Journal of cell biology, 143(2):359–373, 1998.

91. Christos Lamprakis, Ioannis Andreadelis, John Manchester, Camilo Velez-Vega, José S. Duca, and Zoe Cournia. Evaluating the efficiency of the martini force field to study protein dimerization in aqueous and membrane environments. Journal of Chemical Theory and Computation, 17(5):3088–3102, 2021.

92. Mariano A. Ostuni, Patricia Hermand, Emeline Saindoy, Noëlline Guillou, Julie Guellec, Audrey Coens, Claude Hattab, Elodie Desuzinges-Mandon, Anass Jawhari, Soria Iatmanen-Harbi, Olivier Lequin, Patrick Fuchs, Jean-Jacques Lacapere, Christophe Combadìere, Frédéric Pincet, and Philippe Deterre. Cx3cl1 homo-oligomerization drives cell-to-cell adherence. Scientific Reports, 10(1):9069, Jun 2020.

93. Verity Jackson, Julia Hermann, Christopher J Tynan, Daniel J Rolfe, Robin A Corey, Anna L Duncan, Maxime Noriega, Amy Chu, Antreas C Kalli, E Yvonne Jones, et al. The guidance and adhesion protein flrt2 dimerizes in cis via dual small-x3-small transmembrane motifs. Structure, 30(9):1354–1365, 2022.

94. Azadeh Alavizargar, Annegret Elting, Roland Wedlich-Soldner, and Andreas Heuer. Lipid-mediated association of the slg1 transmembrane domains in yeast plasma membranes. The Journal of Physical Chemistry B, 126(17):3240–3256, 2022.

95. Mariana Valério, Diogo A. Mendoņca, Jõao Morais, Carolina C. Buga, Carlos H. Cruz, Miguel A.R.B. Castanho, Manuel N. Melo, Cláudio M. Soares, Ana Salomé Veiga, and Diana Lousa. Parainfluenza fusion peptide promotes membrane fusion by assembling into oligomeric porelike structures. ACS Chemical Biology, 17(7):1831–1843, 2022. PMID: 35500279.

96. J. Karl Spinti, Fernando Neiva Nunes, and Manuel N. Melo. Room for improvement in the initial martini 3 parameterization of peptide interactions. Chemical Physics Letters, 819:140436, 2023.

97. Ainara Claveras Cabezudo, Christina Athanasiou, Alexandros Tsengenes, and Rebecca C. Wade. Scaling protein–water interactions in the martini 3 coarse-grained force field to simulate transmembrane helix dimers in different lipid environments. Journal of Chemical Theory and Computation, 19(7):2109–2119, 2023. PMID: 36821400.

98. Herre Jelger Risselada, Carsten Kutzner, and Helmut Grubmüller. Caught in the act: Visualization of snare-mediated fusion events in molecular detail. ChemBioChem, 12(7):1049–1055, 2011.

99. Hiromi Sesaki and Robert E. Jensen. UGO1 Encodes an Outer Membrane Protein Required for Mitochondrial Fusion. Journal of Cell Biology, 152(6):1123–1134, 03 2001.

100. Hiromi Sesaki and Robert E. Jensen. Ugo1p links the fzo1p and mgm1p gtpases for mitochondrial fusion*. Journal of Biological Chemistry, 279(27):28298–28303, 2004.

101. Fabian Anton, Julia M. Fres, Astrid Schauss, Benôıt Pinson, Gerrit J. K. Praefcke, Thomas Langer, and Mafalda Escobar-Henriques. Ugo1 and Mdm30 act sequentially during Fzo1-mediated mitochondrial outer membrane fusion. Journal of Cell Science, 124(7):1126–1135, 04 2011.

102. Leonid V Chernomordik and Michael M Kozlov. Mechanics of membrane fusion. Nat Struct Mol Biol, 15(7):675–83, 2008.

103. Holger A. Scheidt, Katja Kolocaj, David B. Konrad, James A. Frank, Dirk Trauner, Dieter Langosch, and Daniel Huster. Light-induced lipid mixing implies a causal role of lipid splay in membrane fusion. Biochimica et Biophysica Acta (BBA) - Biomembranes, 1862(11):183438, 2020.

104. Lydie Vamparys, Romain Gautier, Stefano Vanni, W. F. Drew Bennett, D. Peter Tieleman, Bruno Antonny, Catherine Etchebest, and Patrick F. J. Fuchs. Conical lipids in flat bilayers induce packing defects similar to that induced by positive curvature. 104(3):585–593.

105. Stefano Vanni, Hisaaki Hirose, Héléne Barelli, Bruno Antonny, and Romain Gautier. A sub-nanometre view of how membrane curvature and composition modulate lipid packing and protein recruitment. Nature Communications, 5(1):4916, Sep 2014.

106. D. Langosch, M. Hofmann, and C. Ungermann. The role of transmembrane domains in membrane fusion. Cellular and Molecular Life Sciences, 64(7):850, Feb 2007.

107. Mohamed M Elsutohy, Veeren M Chauhan, Robert Markus, Mohammed Aref Kyyaly, Saul JB Tendler, and Jonathan W Aylott. Real-time measurement of the intracellular ph of yeast cells during glucose metabolism using ratiometric fluorescent nanosensors. Nanoscale, 9(18):5904– 5911, 2017.

108. Shailendra S. Rathore, Yinghui Liu, Haijia Yu, Chun Wan, MyeongSeon Lee, Qian Yin, Michael H. B. Stowell, and Jingshi Shen. Intracellular Vesicle Fusion Requires a Membrane-Destabilizing Peptide Located at the Juxtamembrane Region of the v-SNARE. Cell Reports, 29(13):4583– 4592.e3, December 2019.

